# TRIM7 ubiquitinates SARS-CoV-2 membrane protein to limit apoptosis and viral replication

**DOI:** 10.1101/2024.06.17.599107

**Authors:** Maria Gonzalez-Orozco, Hsiang-chi Tseng, Adam Hage, Hongjie Xia, Padmanava Behera, Kazi Afreen, Yoatzin Peñaflor-Tellez, Maria I. Giraldo, Matthew Huante, Lucinda Puebla-Clark, Sarah van Tol, Abby Odle, Matthew Crown, Natalia Teruel, Thomas R Shelite, Vineet Menachery, Mark Endsley, Janice J. Endsley, Rafael J. Najmanovich, Matthew Bashton, Robin Stephens, Pei-Yong Shi, Xuping Xie, Alexander N. Freiberg, Ricardo Rajsbaum

## Abstract

SARS-CoV-2 is a highly transmissible virus that causes COVID-19 disease. Mechanisms of viral pathogenesis include excessive inflammation and viral-induced cell death, resulting in tissue damage. We identified the host E3-ubiquitin ligase TRIM7 as an inhibitor of apoptosis and SARS-CoV-2 replication via ubiquitination of the viral membrane (M) protein. *Trim7^-/-^* mice exhibited increased pathology and virus titers associated with epithelial apoptosis and dysregulated immune responses. Mechanistically, TRIM7 ubiquitinates M on K14, which protects cells from cell death. Longitudinal SARS-CoV-2 sequence analysis from infected patients revealed that mutations on M-K14 appeared in circulating variants during the pandemic. The relevance of these mutations was tested in a mouse model. A recombinant M- K14/K15R virus showed reduced viral replication, consistent with the role of K15 in virus assembly, and increased levels of apoptosis associated with the loss of ubiquitination on K14. TRIM7 antiviral activity requires caspase-6 inhibition, linking apoptosis with viral replication and pathology.

## INTRODUCTION

The severe acute respiratory syndrome coronavirus 2 (SARS-CoV-2) is a highly transmissible positive single-stranded RNA virus of the *Coronaviridae* family ^1,2^. Its RNA genome encodes four structural proteins, which include Spike (S), Nucleocapsid (N), Envelope (E), and Membrane (M) proteins ^3^. Of the structural proteins, M is the most abundant in the virion and is essential for the sorting of structural proteins to promote assembly and release of viral particles ^4^.

The pathogenesis of SARS-CoV-2 in humans includes a combination of excessive inflammatory responses and viral-induced tissue damage that causes lung injury, called acute respiratory distress syndrome ^5^ ^2,6,7^. The severity of disease and respiratory failure in human-infected patients correlates with the increased presence of cytokines including IL-1-β, IL-6, and TNF-α in serum and BAL ^8,9^. SARS-CoV-2 infects multiciliated cells of the respiratory tract and alveolar type 2 (AT2) cells expressing the ACE2 receptor and the TMPRSS2 protease ^5,10–12^. Viral infection can increase levels of apoptosis and other forms of cell death leading to tissue damage. Together, increased cell death and enhanced inflammation can correlate with disease ^12–14^. Multiple mechanisms have been proposed to promote cell death during infection, including cytokine-induced or intrinsic apoptosis directly triggered by viral proteins, including M ^15,16^. The M protein can also promote apoptosis by inhibiting the activation of the PDK1- AKT pathway ^17^, or by inducing mitochondrial intrinsic apoptosis^18^.

The innate immune response elicited by SARS-CoV-2 infection includes innate lymphoid cells, monocytes, macrophages, and neutrophils ^19,20^. In COVID-19 patients, low levels of circulating lymphocytes and increased levels of neutrophils correlate with the severity of the infection ^21,22^. However, neutrophils are not the main cell type found in the lungs of human patients with prolonged severe disease, and there is evidence suggesting that neutrophils may be protective early during infection ^19^. It is still unclear which specific factors during SARS-CoV-2 infection contribute to this shift from protective to detrimental responses by neutrophils and monocytes. Similarly, type-I interferons (IFN-I), which are well-known antiviral cytokines, can play protective or detrimental roles during infection depending on timing ^23,24^, however, it is also unclear what factors determine protective IFN-I induction.

The innate antiviral response against SARS-CoV-2 is mediated primarily by the cytosolic nucleic acid sensor melanoma differentiation-associated protein 5 (MDA5) that recognizes the viral RNA to activate the mitochondrial antiviral signaling protein (MAVS) leading to the production of IFNs and proinflammatory cytokines ^25,26^. Multiple viral proteins, including M, can inhibit the IFN-I pathway by targeting cytosolic receptors or by promoting the degradation of the TANK-binding kinase (TBK-1), reducing IRF3 phosphorylation and IFN-I induction ^27–33^.

TRIM7 belongs to a large family of E3-Ubiquitin (Ub) ligases, which transfer Ub to target proteins ^34^ and can play protective or detrimental roles during infection. TRIM7 has been reported to play antiviral roles against Enteroviruses ^35,36^, and proviral roles during Zika virus infection (ZIKV) ^37^. TRIM7 can also regulate immune responses by promoting the production of IFN-β, TNF-α, and IL-6 in macrophages after TLR4 stimulation ^38^. Although there is previous evidence that TRIM7 may interact with M ^39^ and previous reports identified another E3-ligase, RNF5, as a proviral factor by ubiquitinating M on

K15 ^40^, the pathophysiological roles of TRIM7 and ubiquitination of M *in vivo* during SARS-CoV-2 infection remains unknown. Here we characterized in detail the multiple roles of TRIM7 during infection *in vivo*. We found that TRIM7 regulates the expression of inflammatory cytokines, including the chemokine CXCL1, which promotes the recruitment of immune cells to the infection site. TRIM7 also acts as an antiviral factor during SARS-CoV-2 infection, by ubiquitinating the M protein and inhibiting caspase-6- dependent apoptosis, in an IFN-I independent manner. We also identified the presence of natural K14 mutations in circulating SARS-CoV-2 during the pandemic, supporting a physiological role for ubiquitination of M.

## RESULTS

### TRIM7 Ubiquitinates SARS-CoV-2 M protein on the K14 residue

Previous mass spectrometry studies identified TRIM7 as a potential binding partner of SARS-CoV-2 M protein ^39^, although this interaction and its functional relevance were not further investigated. We first confirmed that TRIM7 and M interact using co-immunoprecipitation assays (coIP) (Fig.1a-b), and this interaction is mediated by TRIM7’s PRY-SPRY C-terminal domain (Fig. S1a). As previously proposed ^39,40^, a significant proportion of M is ubiquitinated when ectopically expressed (Fig. S1b). In addition, overexpression of TRIM7 further enhanced the ubiquitination of M (Fig. 1b-c), whereas a catalytically inactive mutant lacking the RING domain of TRIM7 did not (Fig. 1c). Ectopic expression of the ovarian tumor deubiquitinase (OTU), which cleaves endogenous polyubiquitin chains (polyUb) from modified proteins ^37^, removed all polyUb that coimmunoprecipitated with M, while a catalytically inactive mutant of OTU (OTU- 2A) was used as a negative control (Fig. 1c). These results confirm that TRIM7 promotes ubiquitination of SARS-CoV-2 M protein.

**Figure 1.**
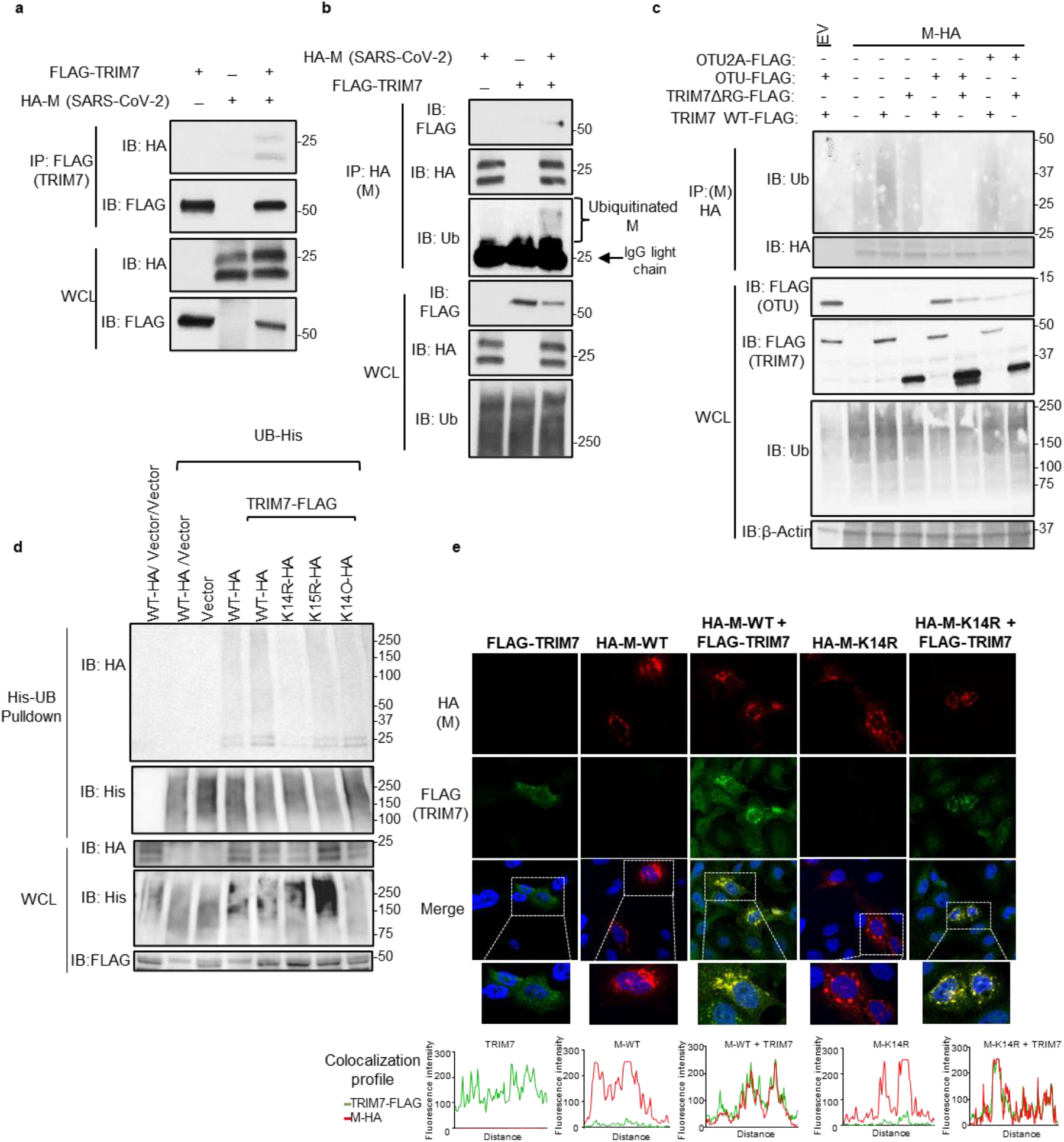
**TRIM7 ubiquitinates SARS-CoV-2 Membrane protein**. a-b) HEK 293T cells were transfected with Vector, M-HA, +/- TRIM7 WT-FLAG, and immunoprecipitated using anti-FLAG beads (a) or anti-HA beads (b). c) HEK293T cells transfected with M- HA, TRIM7 WT-FLAG or TRIM7-ΔRING-FLAG +/- OTU-WT-FLAG or OTU-2A-FLAG followed by immunoprecipitation with anti-HA beads, d) denaturing pulldown using NiNTA beads. HEK293T cells transfected with His-Ub, M WT-HA, M-K14R, M-K15R, M- K14O (all Ks mutated to Rs except for K14), +/- TRIM7 WT-FLAG. e) Confocal microscopy of Hela cells transfected with TRIM7-FLAG (488), M WT-HA or M K14R-HA (555), for 24 hours. Colocalization profile graphs are shown.

We next asked whether TRIM7 ubiquitinates a specific lysine residue on M. A denaturing pulldown of ectopically expressed His-tagged Ub and M encoding K-to-R mutations, showed reduced TRIM7-mediated ubiquitination on an M-K14R mutant as compared to WT-M (Fig. 1d). The K15 residue on M, which is ubiquitinated by the E3- ligase RNF5 ^40^, did not appear to be an acceptor for ubiquitination by TRIM7. In further support of this, a K14-only mutant of M, in which all its K residues were mutated to R except for K14 (K14O), showed similar ubiquitination levels as compared to WT M or the K15R mutant, in the presence of overexpressed TRIM7 (Fig.1d). Together, these data suggest that TRIM7 specifically ubiquitinates the M protein on the K14 residue.

Ubiquitination of M on K15 by RNF5 is necessary to promote the efficient formation of virus-like-particles (VLPs) and virus release ^40^. To rule out a functional role for TRIM7 in virus release, we evaluated the efficiency of VLP formation upon transfection of all viral structural proteins in A549 WT or TRIM7 knockout cells (KO), previously generated in our lab ^37^. No apparent differences were observed in the amount of VLPs released from WT and TRIM7 KO (Fig. S1c), suggesting that TRIM7 does not affect virus release, which is consistent with previous observations that the M-K14R mutant is still able to form VLPs ^40^.

Finally, upon ectopic expression, TRIM7 re-localized from discrete punctate cytoplasmic bodies to larger vesicle-like compartments where it colocalized with M (Fig. 1e). In addition, M localized in the Golgi compartment (Fig. S1d), as in previous reports ^41^. The M-K14R mutant still colocalized and coimmunoprecipitated with TRIM7 (Fig. 1e and S1e), indicating that the interaction did not depend on M-K14 ubiquitination.

Overall, these data provide evidence that TRIM7 specifically ubiquitinates M on its K14 residue, and this ubiquitination does not affect M’s function in the assembly and release of viral particles.

### TRIM7 has antiviral activity during SARS-CoV-2 infection

Since TRIM7 has been reported to have both proviral and antiviral roles, we next evaluated the role of TRIM7 during SARS-CoV-2 infection. Overexpression of TRIM7 in HEK293T cells stably expressing human ACE2 (293T-hACE2) significantly reduced SARS-CoV-2 titers (plaque assay) and viral RNA (qPCR) as compared to the inactive TRIM7-ΔRING, or a vector control (Fig. 2a-b). We then tested whether TRIM7 also has antiviral function *in vivo.* WT and *Trim7^-/-^* mice ^37^ were infected with a mouse-adapted strain of SARS-CoV-2 (CMA3p20) ^42^. *Trim7^-/-^* mice lost significantly more weight at the acute phase of infection and showed slower recovery than WT controls (mixed males and females, Fig. 2c). *Trim7^-/-^* male mice exhibited significantly higher lung viral titers (Fig. 2d) and viral RNA at day 2 and 3 p.i., while we observed smaller differences between females (Fig. S2a-c). The increase in weight loss and viral titers in *Trim7^-/-^* mice correlated with clinical scores (e.g., ruffled fur and/or hunched posture, Fig. S2d), as well as consolidation of the airway at later time points (Fig. S2e-f). TRIM7 antiviral effects were most likely independent of the IFN-I response because the levels of IFN-β mRNA were trending higher in *Trim7^-/-^* and correlated with significantly increased levels of ISG54 and CXCL10, which are well-known ISGs (Fig. 2e-g). In further support of this, IFN-I receptor (IFNAR1) blockade resulted in less weight loss without significantly affecting virus titers as compared to isotype control-treated mice (Fig. 2h-i). As expected, anti-IFNAR1 treated mice showed reduced levels of ISGs (Fig. 2j-k). These results are in line with previous reports showing that IFN-I has a pathogenic effect during SARS-CoV-2 infection, by regulating the infiltration of inflammatory cells but not affecting virus levels ^23,24^.

**Figure 2.**
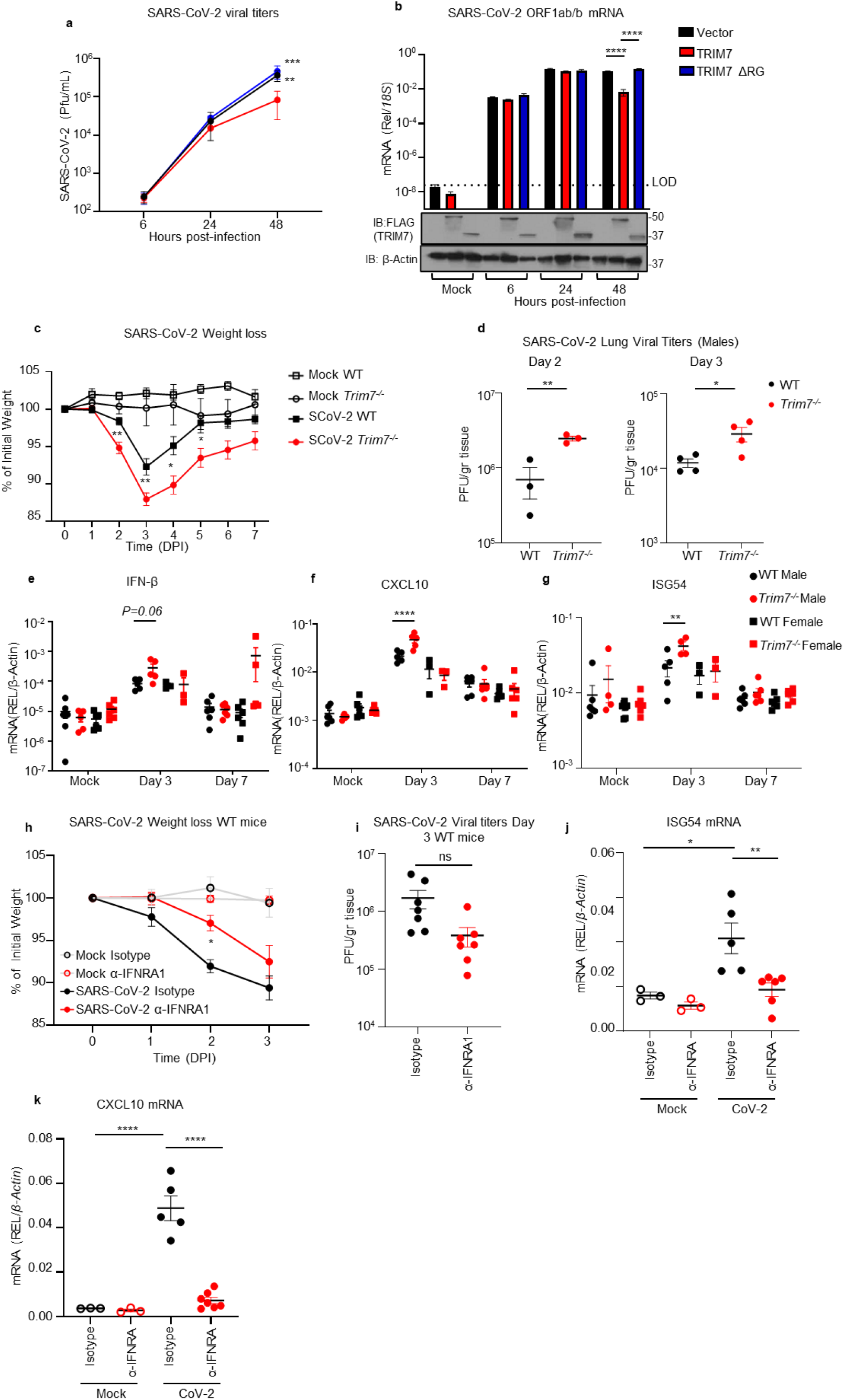
**TRIM7 has antiviral activity during SARS-CoV-2 infection**. a-b) HEK293T- hACE-2 cells were transfected with TRIM7 WT or TRIM7-ΔRING followed by infected with SARS-CoV-2 (MOI 0.1). Viral titers quantified by plaque assay (a) or viral RNA by qPCR (b). Bottom panel in b shows Immunoblot control for TRIM7 expression. c-g) C57BL/6NJ WT (n=18 9 females, 9 males) and *Trim7^-/-^* mice (n=23 10 females, 13 males) infected with maSARS-CoV-2 (1x10^6^ PFU). Weight loss (c), viral titers by plaque assay (d), gene expression in lung tissue by qPCR (e: IFN-β, f: CXCL10, g: ISG54). h-j) IFNAR1 blockade: C57BL/6J WT (n=7/group 5 females and 2 males) were treated with anti-IFNRA1 or isotype 24h before infection with maSARS-CoV-2 (1x10^6^ PFU). Weight loss (h), lung viral titers (i) and gene expression by qPCR of ISG54 (j) CXCL10 (k). Data are depicted as Mean + SEM. (a-b) are representative of 3 independent experiments in triplicates 2-way ANOVA Tukey’s multiple comparisons. (c) is combined data from 3 independent experiments 2-way ANOVA Tukey’s multiple comparisons. (d-g) representative data of 3 independent experiments (d) T-test. (e-g) 2-way ANOVA Tukey’s multiple comparisons. (h) representative data 2-way ANOVA Tukey’s multiple comparisons. (i) T-test analysis. (j-k) representative data one-way ANOVA Tukey’s multiple comparisons. p < 0.001 **, p < 0.0001 ***, p < 0.00001 ****.

Together, our data indicates that TRIM7 plays an antiviral role in cell culture and *in vivo*. These effects require TRIM7 E3-ubiquitin ligase activity and do not appear to be IFN mediated.

### TRIM7 is a negative regulator of IFN-I induction during SARS-CoV-2 infection

Although our data suggest that IFN-I is not involved in the TRIM7-mediated antiviral response, elevated IFN induction could still affect inflammatory responses leading to disease. Therefore, we determined whether the increased levels of ISGs observed in *Trim7^-/-^* infected mice are due to a direct effect of TRIM7 in the IFN pathway. TRIM7 represses expression of IFN-I because bone-marrow derived dendritic cells (BMDCs) from *Trim7^-/-^* mice infected with SARS-CoV-2 showed increased levels of IFN-β mRNA as compared to WT BMDCs (Fig. S2g). In contrast, *Trim7^-/-^* cells expressed lower levels of IL-1β mRNA when compared with WT BMDCs (Fig. S2h). Although SARS-CoV-2 does not productively replicate in DCs ^43^, the presence of similar levels of viral RNA in WT and KO cells indicated that the effects observed are not due to differences in virus infection (Fig. S2i).

To further examine how TRIM7 inhibits IFN-I induction, we tested interactions with the pattern recognition receptors (PRRs) RIG-I and MDA5. Results from coIP assays revealed that TRIM7 interacts with both PRRs (Fig. S2j). Since there is evidence that MDA5 is the major cytosolic receptor for SARS-CoV-2 ^25,26^, we also evaluated if TRIM7 can affect MDA5’s induction of IFN-β. IFN luciferase reporter assays showed that increased concentrations of TRIM7 reduced the IFN-β promoter activity (Fig. S2k), suggesting that TRIM7 can negatively regulates IFN-β by inhibiting MDA5-mediated signaling.

### Ubiquitination on M-K14 does not affect IFN-I antagonist function

M has also been shown to inhibit both the IFN-I production as well as the IFN-I signaling pathways ^27–30,44^. Therefore, we examined if M ubiquitination can affect IFN antagonism. Ectopic expression of WT M or a mutant lacking all ubiquitination sites (M-KallR) inhibited IRF3 phosphorylation at comparable levels upon stimulation with the dsRNA mimic poly (I:C) (Fig. S2l), suggesting that ubiquitination on M does not play a role in inhibition of IFN-I production. M has also been reported to inhibit the induction of ISGs downstream of the IFN-I receptor ^30^. M WT, as well as the mutants M-K14R, M-K15R, and M-KallR reduced the IFN-induced ISRE luciferase reporter activity (Fig. S2m), suggesting that ubiquitination of the lysine residues is not necessary for antagonism of IFN-I signaling.

Taken together, the increased IFN response observed in *Trim7^-/-^* mice is unlikely to be mediated by M ubiquitination and does not explain the increased virus replication observed in the knockout mice.

### TRIM7 promotes innate immune inflammation while protecting from cell death during SARS-CoV-2 infection

TRIM7 has been associated with the induction of genes involved in cell growth, proliferation, and survival ^45^. Conversely, SARS-CoV-2 has been shown to induce cell death in lung cells ^46^, and its M protein has been associated with apoptotic effects ^17^. At day 3 post-infection, we observed a significantly higher proportion of cells positive for Apotracker staining (cells undergoing apoptosis) in the lungs of *Trim7^-/-^* mice as compared to WT controls (Fig. 3a). These effects were evident in CD45^-^ cells (Fig. 3b-c and S3a-c). In contrast, no differences were observed in apoptosis between WT and KO mice within the hematopoietic CD45^+^ compartment (Fig. S3b).

**Figure 3.**
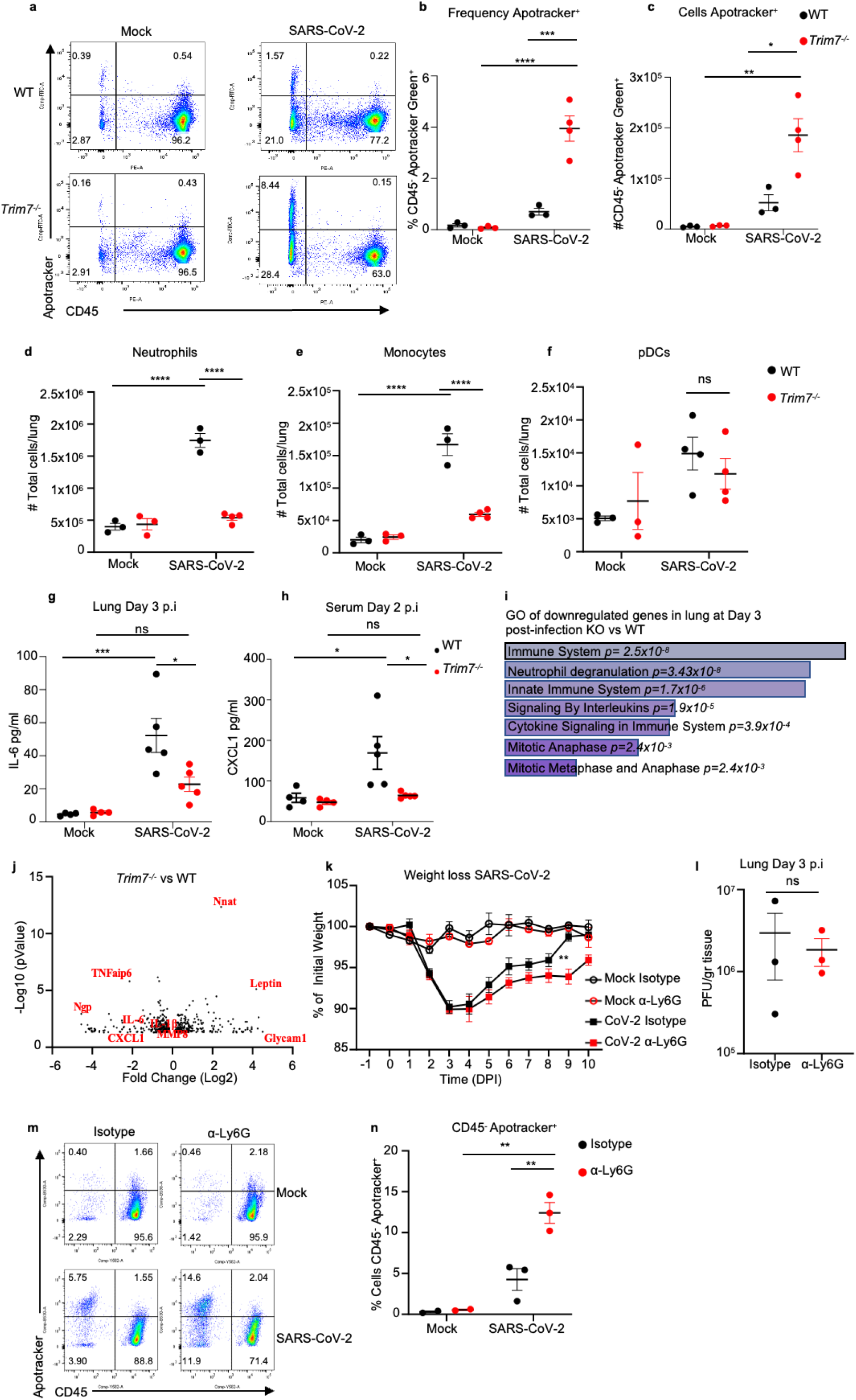
***Trim7 ^-/-^* mice have impaired innate immune response to SARS-CoV-2 infection**. Male C57BL/6NJ WT and *Trim7^-/^*^-^ mice were infected with maSARS-CoV-2 (WT n=3, KO n=4). At day 3 post-infection lung cells were stained for CD45 and Apotracker Green and a representative FACS dot plot shown in (a), frequency of CD45- Apotracker Green^+^ cells (b), or total number of cells per lung (c), total number of neutrophils CD45^+^CD11c^-^CD11b^+^Ly6C^int^Ly6G^hi^ (d), monocytes CD45^+^CD11c^-^ CD11b^+^Ly6C^hi^Ly6G^-^ (e), pDCs CD45^+^CD11c^lo^PDCA-1^+^CD11b^-^ (f). 23-Bioplex analysis from lung (g) or serum (h). IL-6 and CXCL1 are shown. i) Gene ontology graph of genes downregulated in the lung of *Trim7^-/-^* mice at day 3 post-infection. j) Volcano plot of genes changing in lung in WT and *Trim7^-/-^* mice at day 3 post-infection. k) Weight loss graph of WT mice depleted of neutrophils using anti Ly6G antibody or isotype as control infected with maSARS-CoV-2 (n=8). l) viral titers in the lung of mice infected as (k) (n=3), (m) representative dot blot of lung cells stained using CD45 and Apotracker Green, (n) Frequency of cells CD45- Apotracker Green^+^. Data are depicted as Mean ± SEM. (b-h) and (k), representative data of at least 2 independent experiments 2-way ANOVA Tukey’s multiple comparisons. (l), T-test analysis. (n), 2-way ANOVA Tukey’s multiple comparisons. p < 0.001 **, p < 0.0001 ***, p < 0.00001 ****.

We evaluated whether TRIM7 antiviral effects were associated with changes in the innate immune cell composition in the lungs. *Trim7^-/-^* mice showed reduced neutrophil and monocyte infiltration as compared to WT mice at day 3 p.i. (Fig. 3d-e and S3c), whereas no differences in infiltration of plasmacytoid DCs (pDCs) were observed (Fig. 3f). Multiplex analysis of lung and serum cytokines showed reduced pro-inflammatory cytokines IL-6, IL-1β, and IL-1α in *Trim7^-/-^* mice (Fig. 3g and S4a-b). In line with the reduced cellular infiltration to the lung, the neutrophil chemoattractant CXCL1 in serum was lower in *Trim7^-/-^* mice as compared to controls (Fig. 3h). In further support of the role of TRIM7 in promoting immune inflammation, RNAseq and Gene Ontology analysis (GO) of infected lungs showed that downregulated genes were enriched in pathways related to the inflammatory response (neutrophil degranulation, innate immune signaling, and cytokine signaling) as well as cell division/survival (mitotic genes) (Fig. 3i). Specifically, induction of *Il6*, *Il1b*, *Cxcl1*, *Tnfaip6*, and *Mmp8* was reduced in *Trim7*^-/-^ mice (Fig. 3j). These results correlate at the protein level of IL-6 in the lung (Fig. S4c-d) and CXCL1 in the serum.

Since we found a lower number of neutrophils in the lungs of *Trim7^-/-^* mice, we asked whether neutrophil recruitment to the lung could be associated with protection from disease. To test this, C57BL/6J WT mice were depleted of neutrophils (Fig. S4e). Anti- Ly6G-treated mice lost weight at a similar rate as isotype-treated mice until the peak of viral titers (day 3 p.i.). However, neutrophil-depleted mice recovered from infection significantly slower than control mice (Fig. 3k). These effects did not appear to be due to differences in virus replication because control and neutrophil-depleted mice showed similar virus titers (Fig. 3l). These data suggest that neutrophils are not responsible for the antiviral role mediated by TRIM7 but may be involved in tissue repair/healing during the recovery phase. In support of this, neutrophil-depleted mice had a significantly higher frequency of apoptotic cells, specifically in the CD45^-^ compartment (Fig. 3m-n), suggesting that neutrophils are important for the removal of apoptotic cells either directly or indirectly, promoting tissue repair during the recovery phase.

Overall, our data indicate that TRIM7 is antiviral during SARS-CoV-2 infection and suggest that TRIM7 regulates inflammatory immune responses.

### TRIM7 protects from SARS-CoV-2-induced apoptosis and requires an intact K14 residue on M

Apoptosis during viral infection is a process that can either limit virus replication or promote virus dissemination ^47^. To evaluate the relationship between M, TRIM7, and apoptosis, WT and TRIM7 KO A549 cells were transfected with vectors expressing WT or M mutants. Upon transfection of WT M, a significantly higher frequency of apoptotic cells was observed in TRIM7 KO cells as compared to WT cells. These effects required the presence of an intact K14 residue on M because expression of an M-K14R mutant that cannot be ubiquitinated by TRIM7 induced higher frequency of cells in apoptosis in WT cells, and no further difference was observed in TRIM7 KO cells (Fig. 4a-b, controls for expression shown in Fig. S5a). TRIM7 KO cells also display reduced AKT phosphorylation upon stimulation with TNF, while no differences were observed in IKKα/β phosphorylation (Fig. S5b), providing further evidence that TRIM7 is involved in signaling pathways associated with apoptosis and cytokine signaling. Overall, these data suggest that TRIM7 protects from cell apoptosis via ubiquitination on the K14 residue and this potentially reduce virus replication.

**Figure 4.**
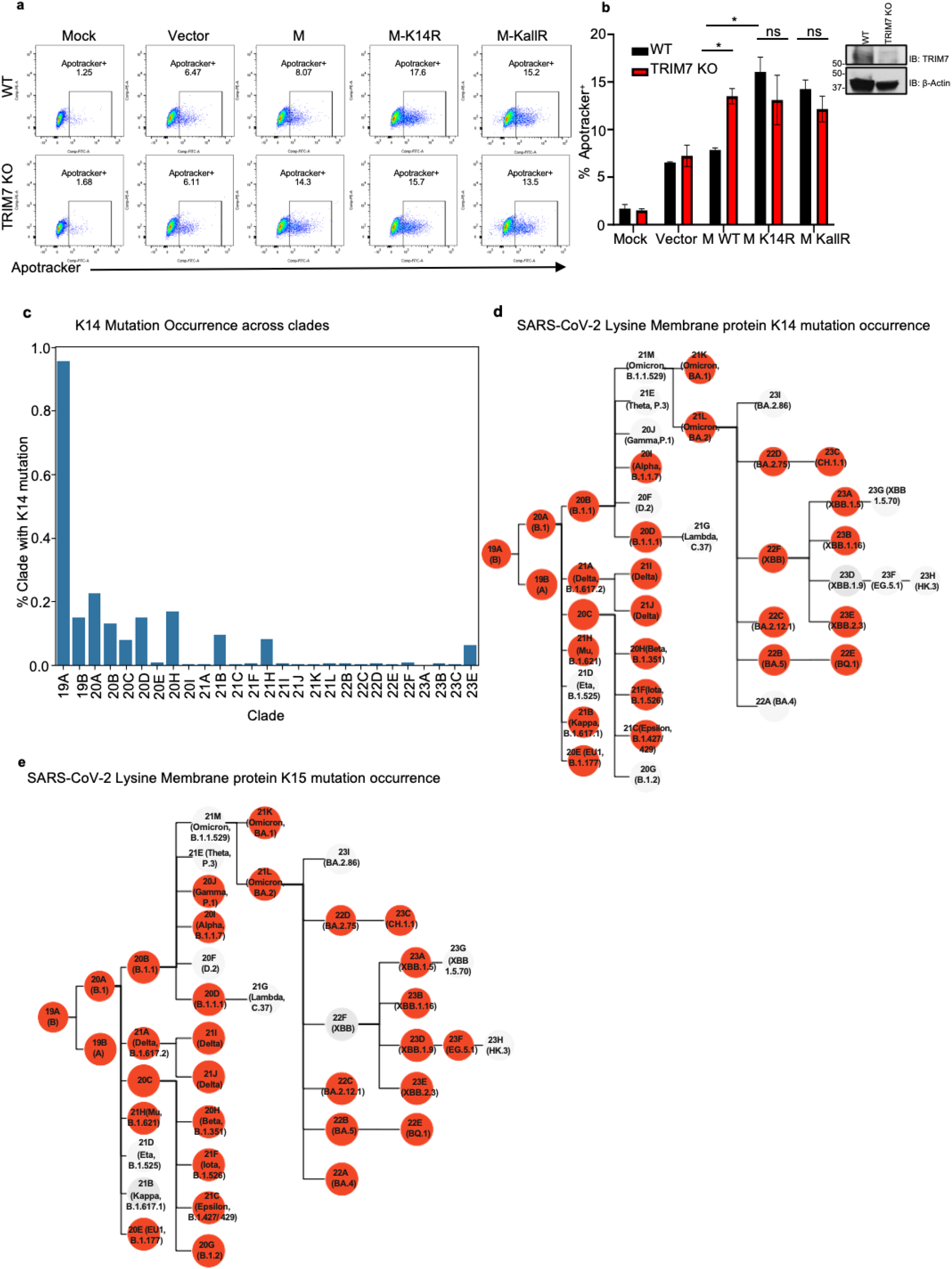
**Mutations on M lysine 14 induce apoptosis and are present in circulating stains of SARS-CoV-2**. a-b) A549 WT or TRIM7 KO transfected with M- WT, M-K14R, or M-KallR mutants for 24h and then stained with Apotracker Green, representative dot blot (a), frequency of cells Apotracker^+^ (b). c-d) M lysine mutations in SARS-CoV-2 sequences. c) percentage of occurrence of M-K14 mutation across the clades. d) membrane protein K14 mutation occurrence. e) membrane protein K15 mutation occurrence. Red highlighted nodes indicate at least one mutation occurrence in the specific clade. Data are depicted as Mean ± SEM. b, representative data of 2 independent experiments in duplicates 2-way ANOVA Tukey’s multiple comparisons test. p < 0.001 **, p < 0.0001 ***, p < 0.00001 ****.

### SARS-CoV-2 Membrane protein mutations on K14 appeared during the pandemic in COVID-19 patients

We next asked whether mutations that lead to loss of ubiquitination and cause more apoptosis can appear in circulating strains of SARS-CoV-2. Data analysis from all ∼8.5 million SARS-CoV-2 genomes present in GenBank from the beginning of the pandemic to March 2024, was performed with CoV-Spectrum ^48^. From the samples analyzed, we observed that 985 showed mutations on K14 residue representing 0.01% of the total samples. Mutations on this site were relatively more frequent early in the pandemic, with a higher occurrence (∼0.95%) in samples from clade 19A (Fig. 4c). The most common mutation was a deletion of K14, followed by the K14R mutation (Fig. 4d and Table 1).

The mutations on the K15 residue were more frequent, present in 0.02% of associated samples in GenBank (2433), Fig. 4e and Table 2). Samples with K14 and K15 mutations were more infrequent, being observed 51 times in total, with all except one of these double mutations being a double deletion (and Table 3). This analysis shows that these mutations can occur in nature and were overrepresented in early clades during the pandemic.

### A recombinant virus with M K14/K15 mutations causes more apoptosis in mice

Since we are unable to correlate these mutations with clinical data from patients, we tested the relevance of these mutations in viral pathogenesis in a mouse model. To this end, we generated a recombinant mouse-adapted double mutant virus M-K14/K15R (CMA5 M-K14/15R), which cannot be ubiquitinated on either K14 or K15 sites. We used this double mutant to avoid generating a virus with increased replication ability due to the loss of the target site for TRIM7 ubiquitination. Since previous studies have shown that ubiquitination on M-K15 by another E3-Ub ligase, RNF5, is required for efficient virus release ^40^, introducing the K15R mutation on the K14R mutant virus should result in an attenuated virus. This would still allow us to dissect the roles of ubiquitination on virus replication and apoptosis by both the K14 and K15 residues of M.

As predicted, this M-K14/K15 mutant virus showed reduced replication kinetics in the IFN-incompetent Vero E6 as well as in IFN-competent Calu-3 cell lines, as compared to the parental WT virus (Fig. 5a, 5c, S6a and S6c). Consistent with the role of K15 in virus particle formation and budding ^40^, the viral RNA accumulated in the cells at similar rates between the K14/15R and the parental virus strain (Fig. 5b, 5d, S6b, 6d). Since Vero cells do not produce active IFN-I, the differences observed are likely not IFN-I dependent. Consistent with this, no difference in ISG54 mRNA levels was observed between the parental and the mutant virus in Calu-3 cells (Fig. S6e). These data contrast with the phenotype we observed of enhanced virus replication in *Trim7^-/-^* mice but can be explained by the loss of ubiquitination on M-K15 that is required for virus release. Importantly, even though the M-K14/15R mutant virus is highly attenuated, it showed a higher ratio of cells in apoptosis when normalized by PFU (Fig. 5e). Similarly, the M-K14/15R virus replicated to lower levels in the lungs of WT mice (Fig. 5f) but caused increased weight loss (Fig. 5g), and increased apoptosis as compared to the WT parental virus (Fig. 5h and S6f-g). No differences in the production of IFN-β or ISG54 were observed in the lung at day 3 p.i. (Fig. S6h-i). Together, these data and the data described above suggest that the K15 site promotes virus replication while the K14 site protects cells from apoptosis during SARS-CoV-2 infection.

**Figure 5.**
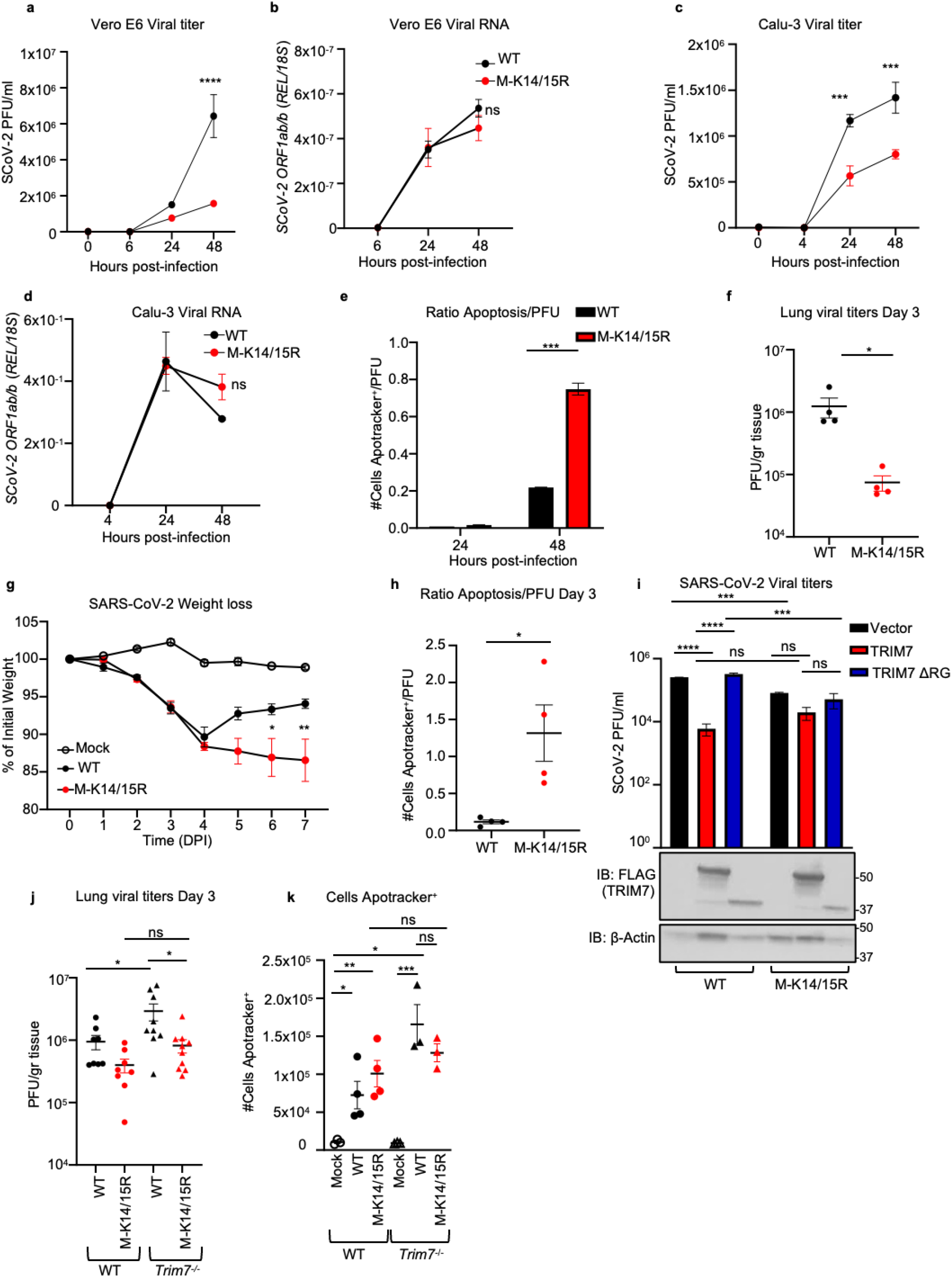
Recombinant virus with M K14/K15 mutations shows increased pathogenesis. Vero E6 or Calu-3 infected with SARS-Cov-2 WT or M-K14/15R MOI 0.1 and 1 respectively. a) viral titers and b) viral RNA in Vero E6. c) viral titers and d, viral RNA in Calu-3. e) ratio of cells in apoptosis Apotracker^+^ normalized by viral titers (Apotracker^+^/PFU) in Calu-3 cells. WT C57BL/6J male mice infected with SARS-CoV-2 WT (n=8) or M-K14/15R mutant viruses (n=10) and mock (n=5). At day 3 post-infection 4 mice were euthanized and lung collected for plaque assay and flow cytometry. f) viral titers in lung. g) weight loss curve. h) ratio of cells in apoptosis in lung (Apotracker^+^/PFU). i) viral titers of HEK 293T-hACE-2 cells transfected with TRIM7 WT or TRIM7ΔRING and infected with SARS-CoV-2 WT or MK14/15R at MOI 0.1 (bottom panel western blot analysis of overexpression of TRIM7). WT (n=8, 6 males, 2 females) and *Trim7^-/-^* (n=9 and 11, 7 or 9 males and 2 females) mice infected intranasal with SARS-CoV-2 WT or M-K14/15R and euthanized at day 3 post-infection. j) lung viral titers. k) total number of cells in apoptosis (CD45 ^+^Apotracker^+^). Data are depicted as Mean ± SEM. a-e) are representative of 3 independent experiments in triplicate 2-way ANOVA Tukey’s multiple comparisons. f-h) is presentative data of at least 2 independent experiments. f) and h) t-test analysis. g) 2-way ANOVA Tukey’s multiple comparisons. i) is presentative data of at least 3 independent experiments in triplicates, 2-way ANOVA Tukey’s multiple comparisons. j) is combined data of 2 independent experiments, k) is representative data of 1 of the 2 independent experiments of j) one-way ANOVA Tukey’s multiple comparisons. p < 0.001 **, p < 0.0001 ***, p < 0.00001 ****.

**Figure 6.**
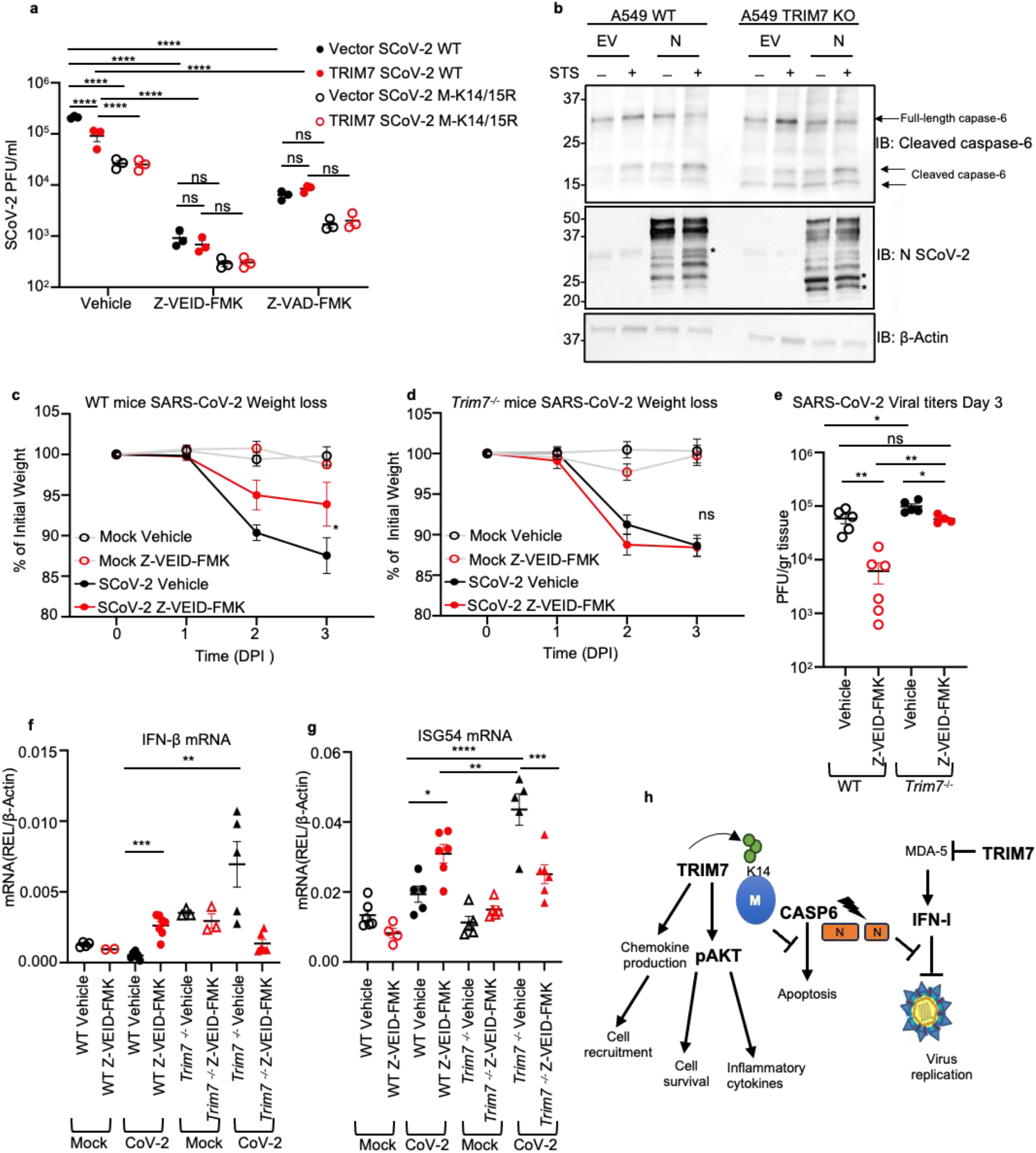
**TRIM7 mediates its antiviral effects by inhibiting caspase-6 activation**. a) viral titers of HEK 293T-hACE-2 cells overexpressing TRIM7 WT and infected with SARS-CoV-2 WT or M-K14/15R at MOI 0.1 and treated with vehicle (DMSO) or 50μM of Z-VEID-FMK 24h post-infection. b) western blot analysis of A549 WT and TRIM7KO cells transfected with SARS-CoV-2 N protein or empty vector and treated with Staurosporine (STS) *indicates cleaved form of N. c) weight loss of WT female mice infected intranasal with SARS-CoV-2 WT treated intraperitoneal with caspase-6 inhibitor or vehicle and lung collected at day 3 post-infection. mocks (n=6 each) vehicle (n=5) or Z-VEID-FMK (n=6) d) weight loss *Trim7^-/-^* female mice as in (c) vehicle mocks (n=5 each) infected vehicle (n=4) or Z-VEID-FMK (n=6). e) lung viral titers. f) IFN-β mRNA and (g) ISG54mRNA expression levels in lung. h) scheme of the multiple functions of TRIM7 during SARS-CoV-2 infection. Data are depicted as Mean ± SEM. a) is presentative data of at least 2 independent experiments in triplicates, 2-way ANOVA Tukey’s multiple comparisons. c-d) are representative data of 2 independent experiments 2-way ANOVA Tukey’s multiple comparisons and e-f) one-way ANOVA Tukey’s multiple comparisons. p < 0.001 **, p < 0.0001 ***, p < 0.00001 ****.

Next, we examined whether TRIM7 can still inhibit virus replication in the absence of the K14/K15 ubiquitination sites. As expected, overexpression of TRIM7 in 293T-hACE2 cells reduced replication of the parental WT virus (Fig. 5i). In contrast, overexpression of TRIM7 did not significantly reduce replication of the K14/15R virus as compared to the empty vector control (Fig. 5i). The mutant virus showed reduced replication as compared to the parental virus, confirming that this mutant virus is attenuated. Overexpression of the inactive TRIM7-ΔRING did not affect the replication of either virus and served as an additional control (Fig. 5i). In line with these results, while the parental WT virus replicated to higher levels in *Trim7^-/-^* compared to WT mice, no significant difference was observed when comparing M-K14/15R titers between WT and *Trim7^-/-^* mice (Fig. 5j). The loss of the K14 ubiquitination site, which is mediated by TRIM7, would explain the lost difference between WT and KO mice. The K14/15R virus still replicated to a lower titer than the WT virus in *Trim7^-/-^* mice and this can likely be explained by the loss of the K15 ubiquitination site, which is dependent on RNF5 and not TRIM7. The loss of ubiquitination on K14/15 resulted in a slight increased number of cells in the lung undergoing apoptosis as compared to the parental virus in WT mice. As expected, the parental virus promoted greater levels of apoptosis in *Trim7^-/-^* mice, however the M-K14/15R mutant virus did not (Fig. 5k). These data suggest that TRIM7 restricts apoptosis in the lung during SARS-CoV-2 infection and this requires intact K14/15 residues on the M protein.

### TRIM7 mediates its antiviral effects by inhibiting caspase-6 activation

Apoptosis during viral infection is known to play an important role in limiting virus replication ^49,50^. Intriguingly, coronaviruses can take advantage of the apoptosis machinery to promote their replication ^51,52^. Therefore, we evaluated if TRIM7’s antiviral mechanism depends on its ability to inhibit apoptosis. To test this, we used inhibitors of apoptosis, Z-VAD-FMK (a pan caspase inhibitor), and Z-VEID-FMK, (which targets caspase-6 and has been shown to inhibit SARS-CoV-2 replication ^51^). Consistent with this previous study, treatment with caspase-6 inhibitor (Z-VEID-FMK) strongly reduced replication of both the parental and the M-K14/15R viruses in 293T-hACE2 cells (Fig. 6a) and completely inhibited apoptosis (Fig. S6j). However, while overexpression of TRIM7 reduced viral titers in DMSO-treated cells, TRIM7 lost its ability to further reduce SARS-CoV-2 replication as compared to vector control in cells treated with Z-VEID-FMK or Z-VAD-FMK (Fig. 6a and expression controls in Fig. S6k). This suggests that TRIM7 requires, at least in part, an active caspase-6 pathway to exert its antiviral activity. Since it has been shown that caspase-6 can cleave N protein of coronaviruses ^52–54^ and cleaved N inhibits the IFN-I response leading to increased virus replication, we evaluated if TRIM7 is involved in the cleavage of N. Treatment with staurosporine (STS), which activates apoptotic pathways, enhanced cleavage of N as compared to vehicle control in WT A549 cells. These effects were further increased in TRIM7 KO cells, in which additional products of N cleavage were evident (Fig. 6b). These effects correlated with slightly enhanced cleavage of caspase-6 in TRIM7 KO cells (Fig. 6b). In support of these results, WT mice treated with Z-VEID-FMK show less weight loss (Fig. 6c) and significantly reduced viral titers in the lungs as compared to DMSO-treated mice (Fig. 6e). In contrast, Z-VEID-FMK treatment of *Trim7^-/-^* mice did not prevent weight loss (Fig. 6d), indicating that TRIM7 deficient mice are more resistant than WT mice to the protective effects of the inhibitor. Importantly, treatment with Z-VEID-FMK reduced virus titers in the *Trim7^-/-^* mice to the levels observed in the DMSO-treated WT animals (Fig. 6e), suggesting that the antiviral activity of TRIM7 is, in part, mediated by inhibition of caspase-6 activity. Notably, TRIM7 also shows antiviral activity *in vivo* that is independent of caspase-6, because *Trim7^-/-^* mice treated with the inhibitor have significantly higher viral titers compared to WT-treated mice (Fig. 6e).

In line with the proposed role of N cleavage in IFN antagonism, WT mice treated with Z-VEID-FMK showed higher levels of IFN-β and ISG54 mRNA in infected lungs, suggesting that the caspase-6 inhibition indeed results in increased IFN responses that could potentially inhibit virus replication, in WT mice. Surprisingly, these effects did not recapitulate in *Trim7^-/-^* mice. Consistent with our data described above, vehicle control treated *Trim7^-/-^* mice showed higher IFN responses than WT mice, however caspase inhibition in *Trim7^-/-^* did not increase but rather reduced IFN/ISGs (Fig. 6f-g). These data suggest that TRIM7 limits virus replication via a mechanism that partially requires inhibiting caspase-6 activity, but it is mostly independent of IFN-I.

## Discussion

In this study, we show that TRIM7 has antiviral activity against SARS-CoV-2 by ubiquitinating the K14 residue on M, and these effects are associated with reduced apoptosis during infection. Our experiments using caspase inhibitors suggest that the antiviral effects of TRIM7 require an active caspase-6 pathway, linking apoptosis to virus replication and pathology. While our study is in line with a previous report that coronaviruses use apoptosis to replicate ^51^, in our study the effects do not seem to be dependent on IFN-I. Although TRIM7 depletion does result in increased cleavage of the viral protein N as well as increased IFN-I induction, these effects do not lead to reduced virus replication. Furthermore, blocking IFN-I signaling did not change virus titers, further suggesting that in this model IFN-I does not play an antiviral role. In previous studies in mice, IFN-I has been associated with pathology ^24^. Our data agree with these studies, in which IFN-I seems pathogenic and not a major antiviral mechanism. Higher levels of IFN produced by the *Trim7^-/-^* mice do not reduce virus titers to the levels observed in WT mice. Intriguingly, blocking IFN-I signaling in the *Trim7^-/-^* mice, which induces higher IFN responses, does not affect weight loss (Fig. S6l), although it does result in increased virus titer as compared to isotype-treated *Trim7^-/-^* mice (Fig. S6m). This suggests that there is a threshold for IFN-I to have antiviral effects, but without affecting pathology. Therefore, the increased disease phenotype observed in *Trim7^-/-^* mice is IFN-I independent.

At the moment the connection between cleavage of N and the increased virus replication observed in *Trim7^-/-^* mice remains unclear. However, it is clear that TRIM7 and M-K14 are associated with inhibiting the caspase-6 pathway to inhibit virus replication.

Our data also indicate that ubiquitination on M-K14 leads to opposite effects from those of the ubiquitination mediated by RNF5 on M-K15, which has a proviral activity ^40^. We further confirmed the previously proposed proviral role of K15, using a recombinant mutant virus.

Although we cannot completely rule out that ubiquitination on K15 can also contribute to effects on apoptosis, our data suggest that ubiquitination on both residues is not mutually exclusive. Using structures of the M protein in its long and short form, we modelled ubiquitinated forms of M with ubiquitin covalently attached to either K14 and/or K15, our structural modeling analysis suggests that ubiquitination of both lysine residues is energetically possible, either with covalent ubiquitination to one lysine in each M protein monomer, or even with ubiquitination happening in neighboring residues of the same chain (Fig. S7a-d). Our calculations indicate that there is a small energetic advantage for the long form of M, suggesting that ubiquitination may drive the population ensemble of M towards the long form.

The advantage of using this double mutant virus is that we can avoid any compensatory ubiquitination on one residue if the neighboring one is missing. We also show that these mutations do not affect the IFN-I response and it is unlikely that the effects on virus replication are IFN-I mediated.

We observed a dysregulation in the inflammatory response in the *Trim7^-/-^* mice with a reduced number of infiltrating neutrophils and monocytes in the lung. Inflammatory monocytes responsible for producing inflammatory cytokines such as IL-6, TNF-α, and IL-1β are recruited to the lung in patients with COVID-19. These cytokines have been associated with detrimental inflammation but can also have protective roles ^58–60^. In this mouse model, the decreased levels of proinflammatory cytokines in *Trim7^-/-^* mice may correlate with dysfunctional activation of the inflammatory responses associated with severe COVID-19 patients ^59,61–63^. Furthermore, a decrease in monocytes in *Trim7^-/-^* correlated with an increase in viral load, consistent with the finding that reduction of monocyte recruitment in *ccr2^-/-^* mice increases virus in the lungs and also increased IFN-I RNA during infection ^64^.

In addition, neutrophils have been associated with pathology through the induction of Neutrophil Extracellular Traps (NETs)^65–67^. Our data suggest that the reduction of neutrophils in *Trim7^-/-^* mice is not the reason for the increased virus titers but could contribute to the increased apoptosis. Neutrophils appear to play an important protective role in the recovery phase and could be associated with healing and protecting from apoptosis. This is in line with studies showing that neutrophils can be involved in tissue repair by MMP-9, which can regulate activation of PRRs and promote angiogenesis ^68–72^, and is relevant given the degree of damage to blood vessels/endothelialitis in COVID-19 ^73^. These effects could be mediated by a specific subpopulation of neutrophils that needs further characterization, that could also potentially be important for the removal of apoptotic bodies or possibly indirectly by recruiting other cells responsible for this clearance.

Together, our data show that TRIM7 is an important regulator of the innate immune inflammatory response that protects against SARS-CoV-2. TRIM7 also negatively regulates MDA5 signaling, which may help control the detrimental inflammatory effects of IFN-I ^74^ (Fig. S8). Finally, we identified that mutations on residues K14 and K15 can occur in the circulating strains of SARS-CoV-2. Although the presence of these mutations is relatively low and does not correlate with a specific variant of concern (VOC), the presence of these mutations could indicate that these strains are potentially more pathogenic. Therefore, we propose that monitoring mutations on M in infected individuals might predict disease severity if the effects can be correlated with clinical profiles in infected patients.

## Acknowledgements

This work was supported by the US National Institute of Health/National Institute of Allergy and Infectious Diseases (NIH/NIAID) grants. R01AI166668, R01AI155466, and P01AI150585 awarded to R.R. R01AI134907 and Building Interdisciplinary Research Careers in Women’s Health Program (BIRCWH) K12HD052023 awarded to M.I.G.

## Author contributions

MG-O. performed all aspects of this study. H.X. generated mutant virus. H-c.T. and A.O. performed *in vivo* experiments. A.H., P.B., K.A., M.I.G., M.H., L.P-C., Y.P-T. and S.v.T. performed *in vitro* experiments. M.C. and M.B. performed global analysis of sequences.

N.T. and R.J.N. performed the computational modeling analysis. T.R.S. performed histopathological analysis. R.S. V.M., M.E., J.E., P-Y.S., X.X., A.F. provided critical reagents and technical advice. R.R. designed, directed, contributed with data analysis, and obtained funding. M.G-O. and R.R. organized the study and prepared the manuscript. All authors read the manuscript and provided comments.

## Declaration of interest

The authors declare no competing interests.

## Resource availability

Further information and requests for resources and reagents should be directed to and will be fulfilled by the corresponding author, Ricardo Rajsbaum

## Materials availability

Plasmids generated in this study are available upon request from the corresponding author.

## Data availability

Transcriptomic data generated during this study has been deposited with the NCBI Gene Expression Omnibus (GEO) database under: GSE268640 accession number.

## METHODS

### Cell culture

HEK293T (CRL-11268), and A549 (CCL-185) cell lines were purchased from ATCC. Calu-3 2B4 cells were kindly provided by Vineet D. Menachery (The University of Texas Medical Branch at Galveston) ^75^ Vero E6 cells were kindly provided by Pei-Yong Shi (The University of Texas Medical Branch at Galveston). HEK293T-hACE2 cells were kindly provided by Benhur Lee (Mount Sinai) ^76^. A549 TRIM7 KO cells were generated as described by ^37^. All cells were maintained in Dulbecco’s Modified Eagle’s Medium (DMEM) (GIBCO) supplemented with 10% v/v fetal bovine serum (FBS) (HyClone) and 1% v/v penicillin-streptomycin (Corning) in a humidified 5% CO2 incubator at 37°C.

### Viruses

Viruses used in this study were handled under biosafety level 3 (BSL-3) conditions at UTMB facilities in accordance with institutional biosafety approvals. SARS-CoV-2 (USA- WA1/2020) was kindly provided by The World Reference Center of Emerging Viruses and Arboviruses (WRCEVA) (The University of Texas Medical Branch at Galveston), SARS-CoV-2 (CMA3p20) was kindly provided by Vineet D. Menachery (The University of Texas Medical Branch at Galveston) and grown in Vero E6 cells as described by Muruato, et al. 2021 ^42^. SARS-CoV-2 USA-WA1/2020+ D614G was provided by Dr. Pei-Yong Shi (The University of Texas Medical Branch at Galveston).

The infectious cDNA clone icSARS-CoV-2 M-K14/15R was constructed through mutagenesis of a mouse-adapted USA-WA1/2020 SARS-CoV-2 (CMA5 strain) used for in vivo studies ^77,78^ . To generate the CMA5 strain, an adaptive mutation (Spike_Q493H) was identified and engineered into the backbone of the CMA3p20 strain ^42^. The full-length cDNA was assembled via *in vitro* ligation and used as a template for T7 *in vitro* transcription. The full-length viral RNA was electroporated into Vero E6 cells. 48 hours post electroporation, the original P0 virus was harvested and used to infect another flask of Vero E6 cells to produce the P1 virus. The titer of the P1 virus was determined by plaque assay on Vero E6 cells. The viral RNA of P1 virus was extracted and sequenced to confirm the designed mutations using the primers: M-K14R/K15R-F- ACCGTTGAAGAGCTTCGCCGCCTCCTTGAACAATGG and M-K14R/K15R-R CCATTGTTCAAGGAGGCGGCGAAGCTCTTCAACGGT. The P1 virus was used for all the experiments performed in this study. All work following electroporation was performed in a BSL3 laboratory.

### Plasmids

The M-WT, M-K14R, M-K15R, M-KallR, and M-K14O were cloned into pXJ-HA plasmid, Flag-TRIM7 constructs Variant 1 and 2 were purchased from Origene (Rockville, MD), the Flag-OTU and -OTU2A were kindly provided by Adolfo Garcia-Sastre (Mount Sinai), the Ub plasmids have been described before ^79^.

### Transfections

Transient transfections of DNA were performed with TransIT-LT1 (Mirus Bio) for HEK293T cells, and Lipofectamine 3000 (Invitrogen) for A549 cells according to the manufacturer’s guidelines. For lipofectamine transfection media was exchanged 6-8 hrs. All transfections were performed in DMEM 10% v/v FBS without penicillin-streptomycin.

### Cell lysis and co-immunoprecipitation

Cells were harvested in RIPA lysis buffer (50 mM Tris-HCl, pH 8.0, 150 mM NaCl, 1% (v/v) IGEPAL CA-630, 0.5% (w/v) sodium deoxycholate, 0.1% (v/v) SDS, protease inhibitor cocktail ^80^, 5 mM N-ethylmaleimide (Sigma), and 5 mM iodoacetamide (Sigma) as deubiquitinase inhibitors. Cell lysates were clarified by centrifugation at 21,000 x g for 20 min at 4C. 10% of the clarified lysate was added to 2X SDS-PAGE loading buffer containing 2- Mercaptoethanol, heated for 30 min at 37°C, and stored at -20°C as a whole-cell lysate (WCL). The remaining lysate was subjected to immunoprecipitation with anti-FLAG M2 or anti-HA, EZview Red agarose beads (Sigma) overnight at 4°C on a rotating platform. Beads were washed seven times in RIPA buffer (150 or 550 mM NaCl) and the bound proteins were eluted using FLAG or HA peptide respectively, elution was reduced in 2X SDS-PAGE loading buffer containing 2- Mercaptoethanol and incubated for 30 min at 37°C.

### Denaturing pull-down

Cell lysis and WCL collection were performed as above. Lysates were subjected to pull down using nickel-nitrilotriacetic acid (Ni-NTA) beads (Qiagen) overnight at 4 °C on a rotation platform. Beads were washed seven times using denaturing buffer containing 50mM Tris HCl pH8.0 (Sigma), 6M urea, 350 mM NaCl, 0.5%(v/v) IGEPAL CA-630 (Sigma) and 40mM imidazole (Sigma). The proteins were eluted at 4 °C for 30 min, using elution buffer containing 50mM Tris-HCl pH8.0 and 300mM imidazole. Eluted proteins were treated with in 2X SDS-PAGE loading buffer containing 2- Mercaptoethanol and incubated for 30 min at 37°C.

### Confocal Immunofluorescence

HeLa cells were seeded into 6-well plates. After 16h, the cells were transfected with 1µg of M WT or M-K14R-HA tagged with the co-expression of TRIM7-FLAG tagged for 24h. The cells were washed with DPBS 1X, fixed with 4% paraformaldehyde 20’, permeabilized with 0.1% Triton X100 (v/v) in DPBS 1X for 5 minutes, and blocked with 0.5% pork skin gelatin (w/v) in DPBS for 1h. The staining was performed with rabbit anti-HA (Milipore Sigma H6908, 1:100 dilution), anti-FLAG (Sigma-Aldrich F1804, 1:100 dilution in P) overnight at 4°C. The next day, cells were washed with DPBS 1X and incubated with the secondary antibodies anti-mouse Ig Alexa Fluor 488 (Invitrogen A21202) and anti-rabbit Ig Alexa Fluor 555 (Invitrogen A31572) at 1:200 dilution each in DPBS 1X) and washed with DPBS 1X after 2h incubation at RT. DAPI staining (Bio Legend) working solution (1µg/mL in PBS) was added for 5 minutes at RT and washed with PBS before mounting with Merck FluorSave^TM^ reagent. Micrographs were taken with the Leica Stellaris 8 tau-STED Microscope (Leica Microsystems). Microscope parameters and LAS-X software post-processing were set constant for each experiment. Fluorescence intensity values were obtained with ImageJ software (National Institute of Health) and curves were graphed with Graphpad Prism 10 (Graphpad Software, Inc.).

### Virus-like particles (VLPs) generation

VLPs were generated by transfection of the plasmids for expression of S-HA, M-HA, N-FLAG, and E-FLAG, into A549 WT and TRIM7 knockout, briefly; 2X10^5^ cells were seeded into a 6-well plate and transfected using a total of 2 µg of plasmid using Lipofectamine 3000 (Invitrogen, USA) as per the manufacturer’s instruction. The molar ratio for the S, E, M, and N plasmid was 8:8:6:3 as described by ^81^. 70h after transfection the supernatant was collected, and the cells were harvested in RIPA buffer for immunoblotting. The supernatant was clarified by centrifugation 4000 rpm for 10 minutes, then the supernatant was filtered through a 0.45 μM mesh to remove the debris, subsequently, the supernatant was layered over in a 20% sucrose gradient and ultra-centrifugated at 25,000 rpm for 3h at 4°C to pellet down the VLPs and subsequently loaded on to discontinuous, 20–60% sucrose solution and centrifuged at 25,000 rpm for 3 h at 4°C. The opaque band containing the VLPs were collected and analyzed by western blot.

### Western blot

Cell lysates were resolved on 7.5 or 4–15% Mini-PROTEAN and Criterion TGX SDS-PAGE gels and transferred to polyvinylidene difluoride (PVDF) membranes using a Trans-Blot Turbo transfer system (Bio-Rad). Membranes were blocked with 5% (w/v) non-fat dry milk in TBST (TBS with 0.1% (v/v) Tween-20) for 1h and then probed with the indicated primary antibody in 3% (w/v) BSA in TBST at 4°C overnight. Following overnight incubation, membranes were probed with secondary antibodies in 5% (w/v) non-fat dry milk in TBS-T for 1 h at room temperature in a rocking platform: anti-rabbit or anti-mouse IgG-HRP conjugated antibody from sheep (both 1:10,000 NA934 and NA931 GE Healthcare). Proteins were visualized using ECL or SuperSignal West Femto chemiluminescence reagents (Pierce) and detected by autoradiography.

### Mice

All animal experiments were carried out following Institutional Animal Care and Use Committee (IACUC) guidelines and have been approved by the IACUC of the University of Texas Medical Branch at Galveston. Our studies utilized 20- to 25-week-old C57BL/6NJ WT mice (The Jackson Laboratory) that match the *Trim7^-/-^* mice generated as described by ^37^ and 25-week-old C57BL/6J WT mice (The Jackson Laboratory). Mice were maintained under specific pathogen-free conditions in the Animal Resource Center (ARC) facility at UTMB. Animal experiments involving infectious viruses were performed under animal biosafety level 3 (ABSL-3) conditions at UTMB in accordance with institutional biosafety approvals.

### *In vitro* virus infection

HEK293t-ACE-2 overexpressing TRIM7 or TRIM7ΔRING domain were seeded onto 24- well plates at a confluency of 100,000 cells/well and infected with SARS-CoV-2 USA/WA-1 D614G strain MOI 0.1 for 1h, cells were washed once with DPBS 1X and incubated with 6, 24 and 48h after infection. supernatant, RNA, and protein were collected to measure virus titers, gene, and protein expression respectively.

### *In vivo* virus infection

WT and *Trim7 ^-/-^* mice were anesthetized with 5% isoflurane and infected intranasal with SARS-CoV-2 1x10^6^ PFU of CMA3p20 strain, mice were weighed every day for 7 days. Euthanasia was performed at days 2, 3, or 7 post-infection using isoflurane overdose, lungs and serum were collected for downstream analysis. For neutrophil depletion experiments WT C57BL/6J mice were injected intraperitoneally ^82^ with 100μg/mouse of anti-Ly6G or isotype (BioXCell) one day before and one after the infection, mice were infected with SARS-CoV-2 1x10^6^ PFU of CMA3p20 and weighed every day for 10days. At Day 3 post-infection a group of mice was euthanized to perform flow cytometry of lung or peripheral blood to confirm neutrophil depletion. For caspase-6 inhibition experiments C57BL/6NJ mice were treated with Z-VEID-FMK (APExBIO), dose: 12.5mg/kg diluted in PBS or DMSO in PBS as the vehicle through IP injection at day 0, 1, and 2 post-infection. For IFN-I blocking experiments mice were IP injected with or IFNAR1 Isotype IgG control antibody 2mg/per mice at day 0.

### Lung single-cell suspension and flow cytometry

Lungs isolated from infected mice were collected in RPMI 10% v/v FBS 1% v/v penicillin-streptomycin, lungs were rinsed with DPBS cut into small pieces and digested in digestion media containing collagenase D 0.7mg/ml and DNase I 30µg/ml in serum-free RPMI for 30 minutes in a humidified 5% CO2 incubator at 37°C. FBS was added to the digestion media to inactivate the enzymes. Lungs were then passed through 70μm cell strainer to obtain single-cell suspension. Red blood cells were lysed using RBC lysing buffer Hybri-Max (Sigma), cells were counted and 1x10^6^ cells were stained using the following antibodies. Anti-CD45-PE(Biolegend), Anti-Podoplanin PE- DAZZLE594 (Biolegend), Anti-CD24-BUV395 (BD Biosciences) Anti-CD31-BV510, Anti- CD326-BV711 (Biolegend), Anti- MHC-II-BV605 (Biolegend), or Anti- PDCA-1-APC, Anti- CD11b-AF700, Anti- Ly6G-BV780, Anti-CD11c Percp-Cy5.5, Anti- Ly6C-FITC. To measure cell death and apoptosis, cells were stained with Ghost dye-Red780(Tonbo Biosciences) or Fixable viability dye-eFluor506(eBiosciences) and Apotracker Green (Biolegend). After staining samples were fixed using 4% ultrapure formaldehyde diluted in DPBS from 16% methanol-free ultrapure formaldehyde (Thermo Scientific) for 48h. Samples were acquired using LSR II Fortessa and analyzed using FlowJo.

### Plaque Assay

The supernatant of infected cells or lung homogenate was used to measure viral titers. Briefly, confluent monolayers of Vero E6 cells plated in a 12-well plate were infected with virus diluted using DMEM 2% v/v FBS without penicillin-streptomycin, incubated at 37°C for 1h rocking the plate every 15 minutes. Infectious were removed and media was replaced with MEM containing 0.6% v/v tragacanth (Sigma), 5% v/v FBS and 1% v/v penicillin-streptomycin, plates were incubated at 37°C for 2 days in humidified 5% CO2 incubator. Plates were fixed and stained using 10% buffered formalin containing 0.5% (w/v) crystal violet for 30 minutes.

### Histology

The right inferior lobe of the lung was fixed in 10% neutral buffered formalin (HT501128, Sigma, MI) for 7 days. Tissues were cut, paraffinized and H&E stained by the Anatomic

Pathology Laboratory of the Pathology Department of University of Texas Medical Branch. The inflammatory score was calculated by analyzing the presence of peribronchiolar infiltrates (Yes=1, No=0) plus 1-2 Foci of inflammation (1), 2-3 foci of inflammation (2) and 3+ foci of inflammation ^83^.

### IFN-β or ISRE luciferase reporter assay

HEK293T cells were seeded into 24-well plates (50,000 cells/well) and were transfected with 30 ng of IFN-β or 180 ng of ISRE reporter plasmid together with 10 ng of Renilla luciferase plasmid. For IFN-β reporter assay cells were co-transfected with 5 ng of MDA5 and increasing concentrations of TRIM7 20,40 or 80 ng for 24hrs. for ISRE assays cells were co-transfected with 100 ng of M WT or K-R mutants plasmids for 24h and stimulated with 1000IU/ml of IFN-β for 16h. Cells were lysed and luciferase activity was measured using the DualLuciferase reporter assay system (Promega) on a Cytation 5 Multi-Mode Reader (BioTek) according to the manufacturer’s instructions. Values were normalized to Renilla.

### Quantitative reverse-transcription-PCR (qRT-PCR)

Total RNA was isolated using the Direct-zol RNA Miniprep Kit (Zymo Research) following the manufacturer’s instructions. Reverse transcription was performed using the High-Capacity cDNA Reverse Transcription Kit (Applied Biosystems). Real-time qPCR was performed in 384-well plates using iTaq Universal SYBR Green Supermix and a CFX384 Touch Real-Time PCR Detection System (Bio-Rad). Gene expression was normalized to either human 18S or murine β-actin by the comparative CT method (DDCT).

### Cytokine quantification

Cytokines from serum and lung homogenate were quantified using Bio-Plex mouse cytokine 23-plex assay (BioRad) following the manufacturer’s instructions. Serum was diluted 1:3. Samples were analyzed in a Bio-Plex200 Multiplex system (Bio-Rad).

### Global Analysis

Global membrane protein K14/K15 mutation occurrence was analyzed using the CoV- Spectrum ^48^ (https://open.cov-spectrum.org) dashboard. The analysis covered all samples in the “Open Data” version of CoV-Spectrum (GenBank deposited samples) and the period 2020-01-06 to 2024-01-31. The following queries were used to determine the occurrence of mutations to K14/K15 and K14+K15 (using Nextstrain Clade 21A as an example): 21A (Nextstrain clade) & M:K14 21A (Nextstrain clade) & M:K15 21A (Nextstrain clade) & M:K14 & M:K15

For each query, the total number of samples belonging to the underlying clade was obtained, and the percentage of samples with the particular mutation was determined using the “Substitutions and Deletions” section of the resulting CoV-Spectrum reports. Data was visualized using an adaption of the nCoV Clades Schema (https://github.com/nextstrain/ncov-clades-schema)^84^ using Miro.

### Methods for M protein + ubiquitin modeling

We utilized structures of the M protein in its long form (PDB 7VGR) and short form (PDB 7VGS). To model the membrane, we used a lipid composition reminiscent of the biological ER-Golgi intermediate compartment (ERGIC) ^55^ and employed the CHARMM-GUI membrane builder ^85–87^. Subsequently, we generated four models of covalent interactions with a ubiquitin structure (PDB 2JF5), each representing interactions with K14 and K15 from either the same M protein monomer or different monomers, for both conformational states of M. These models were then subjected to minimization using the Yasara web server ^56^, followed by the calculation of the total energy of the systems utilizing adapted Surfaces functions ^57^.

## Notes

### Competing Interest Statement

The authors have declared no competing interest.

