## Supplementary data for "TRIM7 ubiquitinates SARS-CoV-2 membrane protein to limit apoptosis and viral replication"

### Author affiliations and footnotes

[1]Department of Microbiology and Immunology, University of Texas Medical Branch, Galveston, TX

[2]Center for Virus-Host-Innate-Immunity, RBHS Institute for Infectious and Inflammatory Diseases, and Department of Medicine, New Jersey Medical School, Rutgers University, Newark, NJ

[3]Department of Biochemistry and Molecular Biology, University of Texas Medical Branch, Galveston, TX

[4]Department of Internal Medicine, Division of Infectious Diseases, University of Texas Medical Branch, Galveston, TX

[5]Hub for Biotechnology in the Built Environment, Department of Applied Sciences, Faculty of Health and Life Sciences, Northumbria University, Newcastle, UK

[6]Department of Pharmacology and Physiology, Faculty of Medicine, Université de Montréal, Montreal, Canada

[7]Center for Immunity and Inflammation and Department of Pharmacology, Physiology and Neuroscience, New Jersey Medical School, Rutgers University, Newark, NJ

[8]Department of Pathology, University of Texas Medical Branch, Galveston, TX

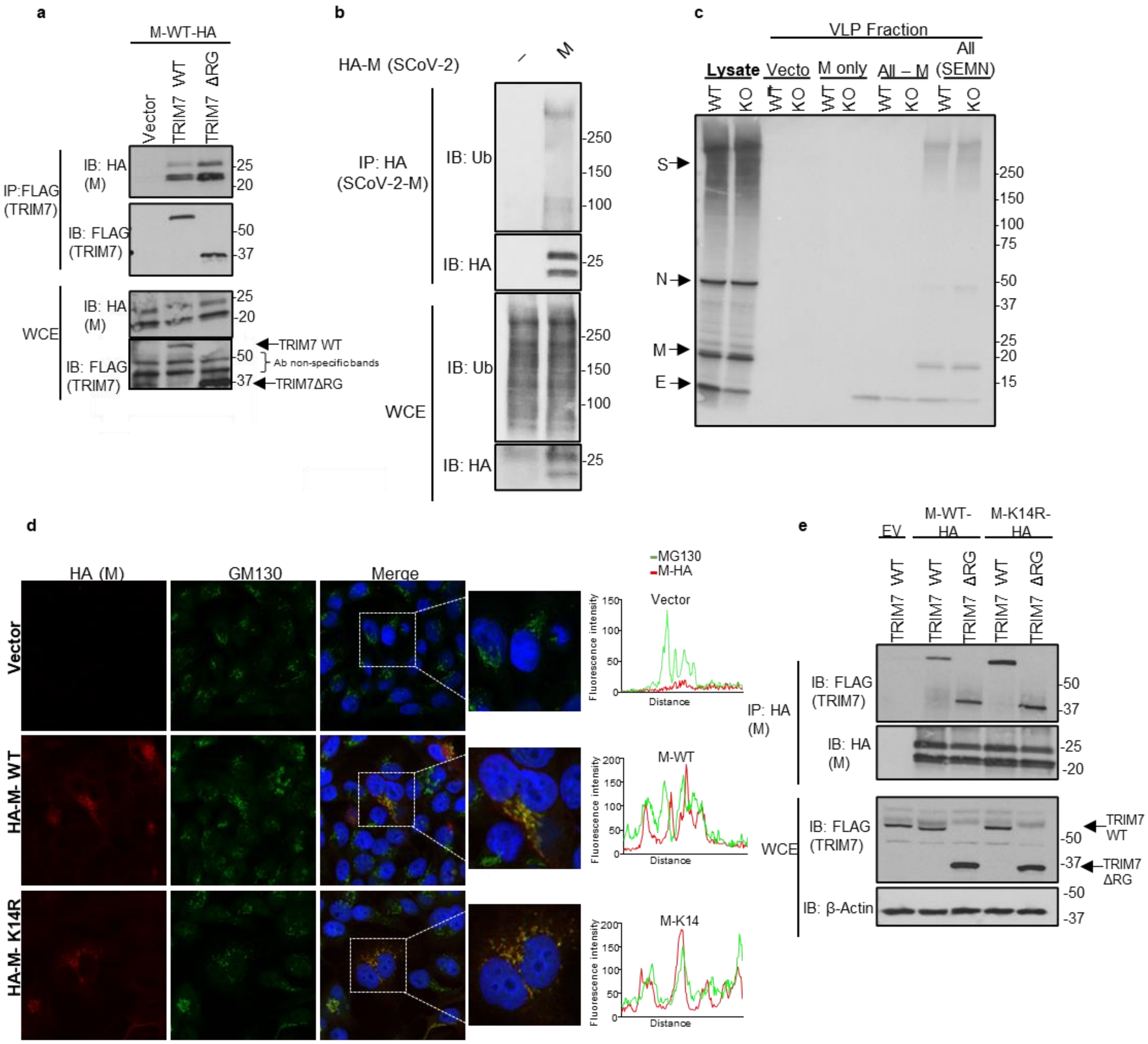


**Figure S1. Ubiquitination of M by TRIM7 does not affect budding.** HEK 293T cells transfected with a) 200ng or TRIM7 WT or ΔRG-FLAG and 100ng of M-HA and immunoprecipitated with FLAG beads. b) 200ng of M-HA and immunoprecipitated using HA beads c) VLPs in supernatant released by A549 WT and TRIM7 KO cells (KO). d, Confocal microscopy of Hela cells expressing M-WT-HA or M-K14R-HA (Red) and label with anti-MG130 (Golgi Marker) for 24 hours. Colocalization profile bottom show the fluorescence intensity. e) 200ng or TRIM7 WT or ΔRG-FLAG and 100ng of M-HA (WT or K14R) and immunoprecipitated with HA beads. Representative experiments of at least 2 independent experiments

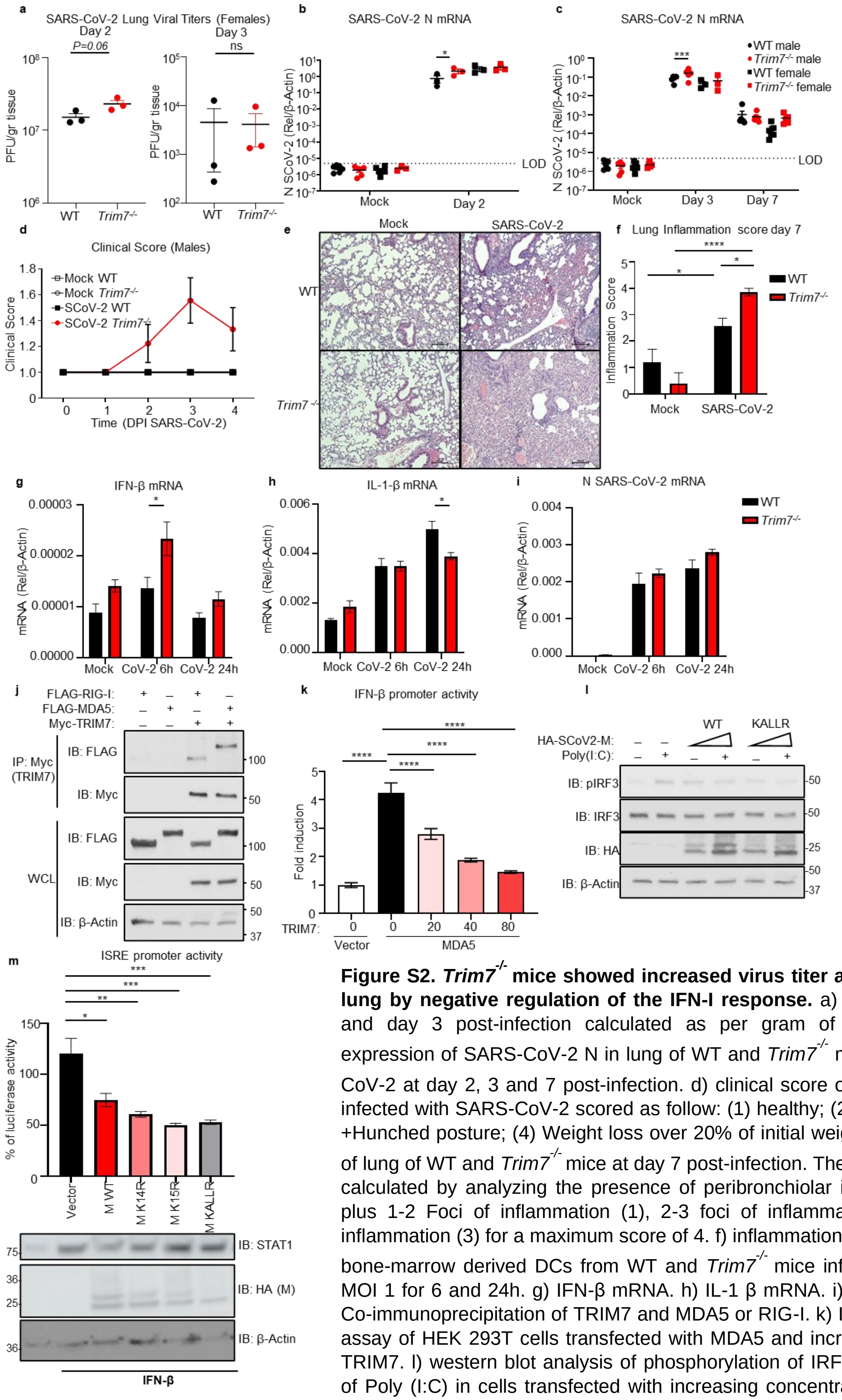


**Figure S2. *Trim7*$^{-/-}$ mice showed increased virus titer and inflammation in the lung by negative regulation of the IFN-I response.** a) lung viral titers at day 2 and day 3 post-infection calculated as per gram of tissue. b-c) viral RNA expression of SARS-CoV-2 N in lung of WT and *Trim7*$^{-/-}$ mice infected with SARS-CoV-2 at day 2, 3 and 7 post-infection. d) clinical score of WT and *Trim7*$^{-/-}$ males infected with SARS-CoV-2 scored as follow: (1) healthy; (2) Ruffled fur; (3) score 2 +Hunched posture; (4) Weight loss over 20% of initial weight. e) histology analysis of lung of WT and *Trim7*$^{-/-}$ mice at day 7 post-infection. The inflammatory score was calculated by analyzing the presence of peribronchiolar infiltrates (Yes=1, No=0) plus 1-2 Foci of inflammation (1), 2-3 foci of inflammation (2) and 3+ foci of inflammation (3) for a maximum score of 4. f) inflammation score. qPCR analysis of bone-marrow derived DCs from WT and *Trim7*$^{-/-}$ mice infected with SARS-CoV-2 MOI 1 for 6 and 24h. g) IFN-β mRNA. h) IL-1 β mRNA. i) N SARS-CoV-2 RNA. j) Co-immunoprecipitation of TRIM7 and MDA5 or RIG-I. k) IFN-β Luciferase reporter assay of HEK 293T cells transfected with MDA5 and increasing concentrations of TRIM7. l) western blot analysis of phosphorylation of IRF3 induced by stimulation of Poly (I:C) in cells transfected with increasing concentrations of M-WT or KallR mutant. m) ISRE luciferase reporter assay of cells transfected with M-WT or mutants K14R, K15R or KallR stimulated with IFN- β for 24h. Data are depicted as Mean ±SEM. T-test analysis, one-way or 2-way Tukey's multiple comparisons tests. $p < 0.001$ **, $p < 0.0001$ ***, $p < 0.00001$ ****.

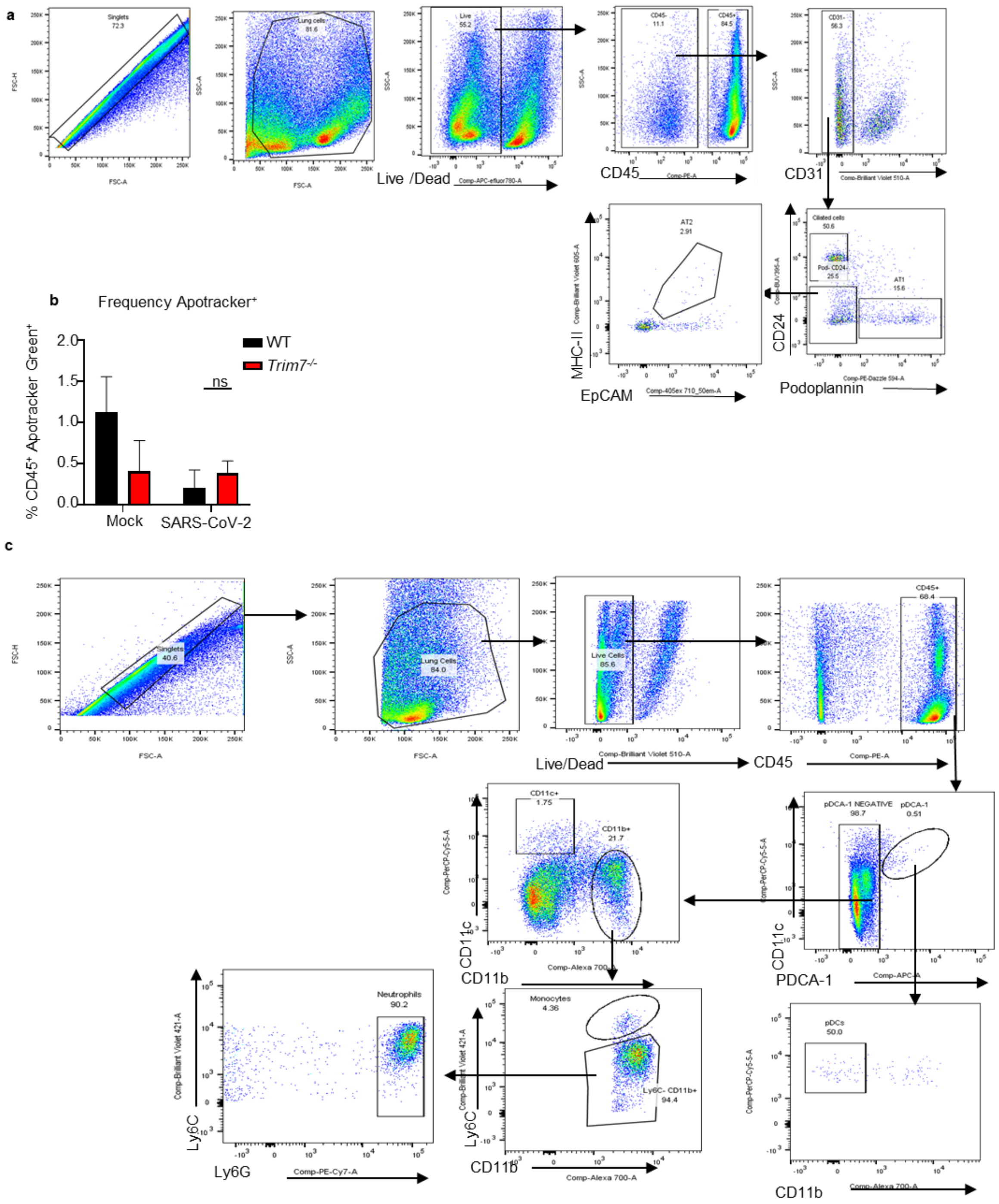


**Figure S3. Flow cytometry gating strategy**. a) flow cytometry gating strategy for lung epithelial cells. b) frequency of cells CD45$^{+}$ Apotracker$^{+}$ in lung of mice infected with SARS-CoV-2 at day 3 post-infection. c) gating strategy for flow cytometry analysis of innate immune cells.

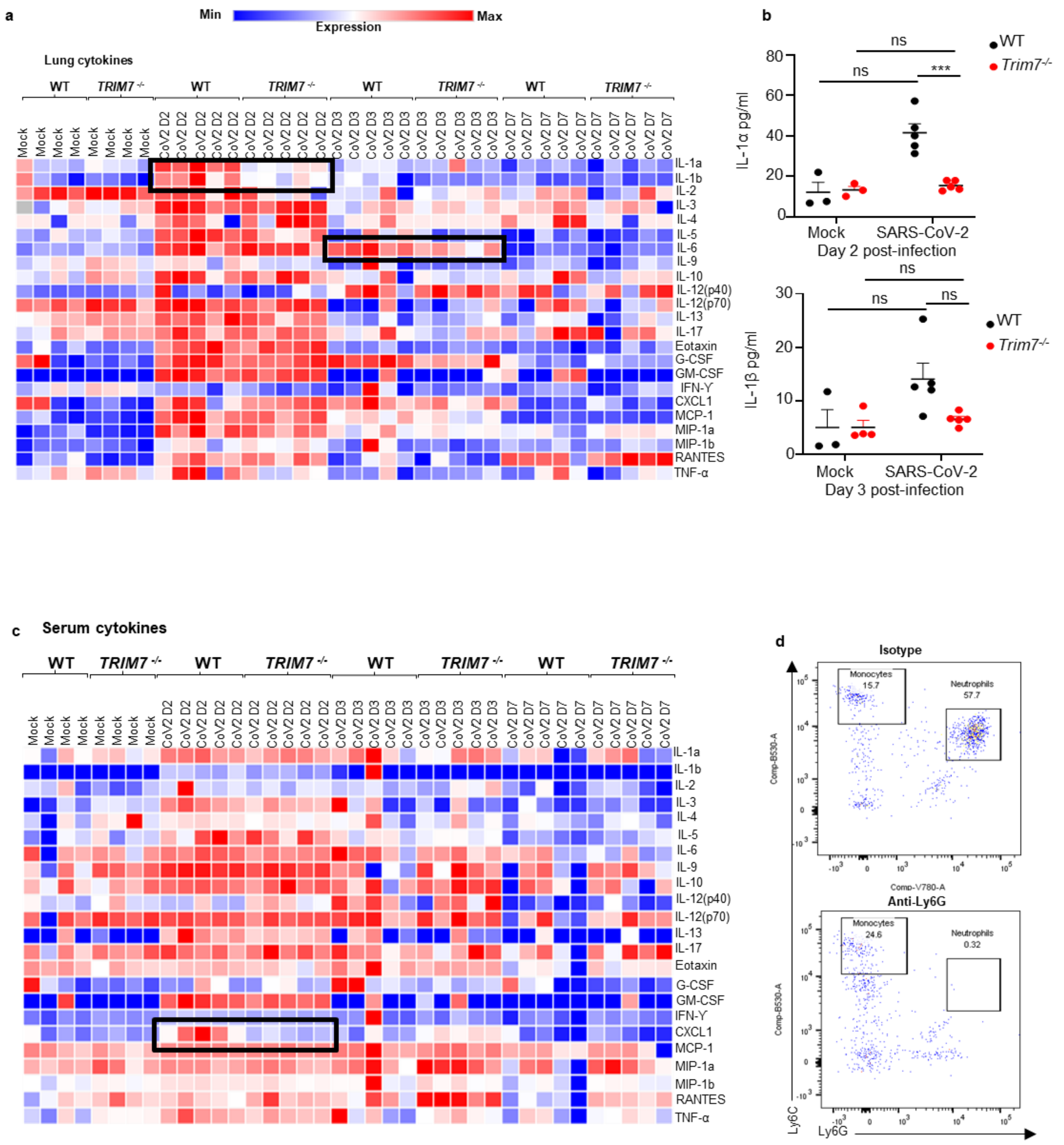


**Figure S4. Multiplex analysis of lung homogenates and serum**. a) heat map of cytokine expression in lung of infected mice. b) lung homogenate concentration levels ofIL-1α in lung of WT and *Trim7*$^{-/-}$ mice. c) heat map of cytokine expression in serum of infected mice. d) representative dot blot of peripheral blood staining showing depletion of neutrophils CD45$^{+}$ CD11c$^{-}$CD11b$^{+}$Ly6C$^{int}$, Ly6G$^{hi}$. Data are depicted as Mean ± SEM. 2-way Tukey's multiple comparisons tests. $p < 0.001$ **, $p < 0.0001$ ***, $p < 0.00001$ ****.

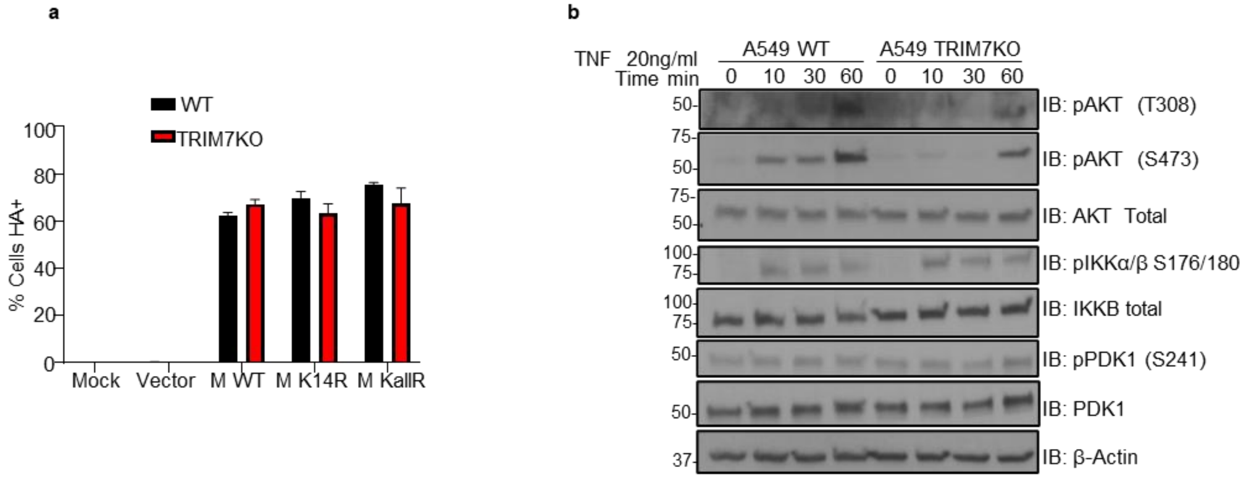


**Figure S5. TRIM7KO cells have impaired activation of AKT pathway after TNF stimulation.** a) frequency of cells transfected with M-WT, M-K14R and M-KallR. b) western blot of A549 WT and TRIM7 KO starved for 8h and then stimulated with 20ng/ml of TNF for 10, 30 and 60 minutes.

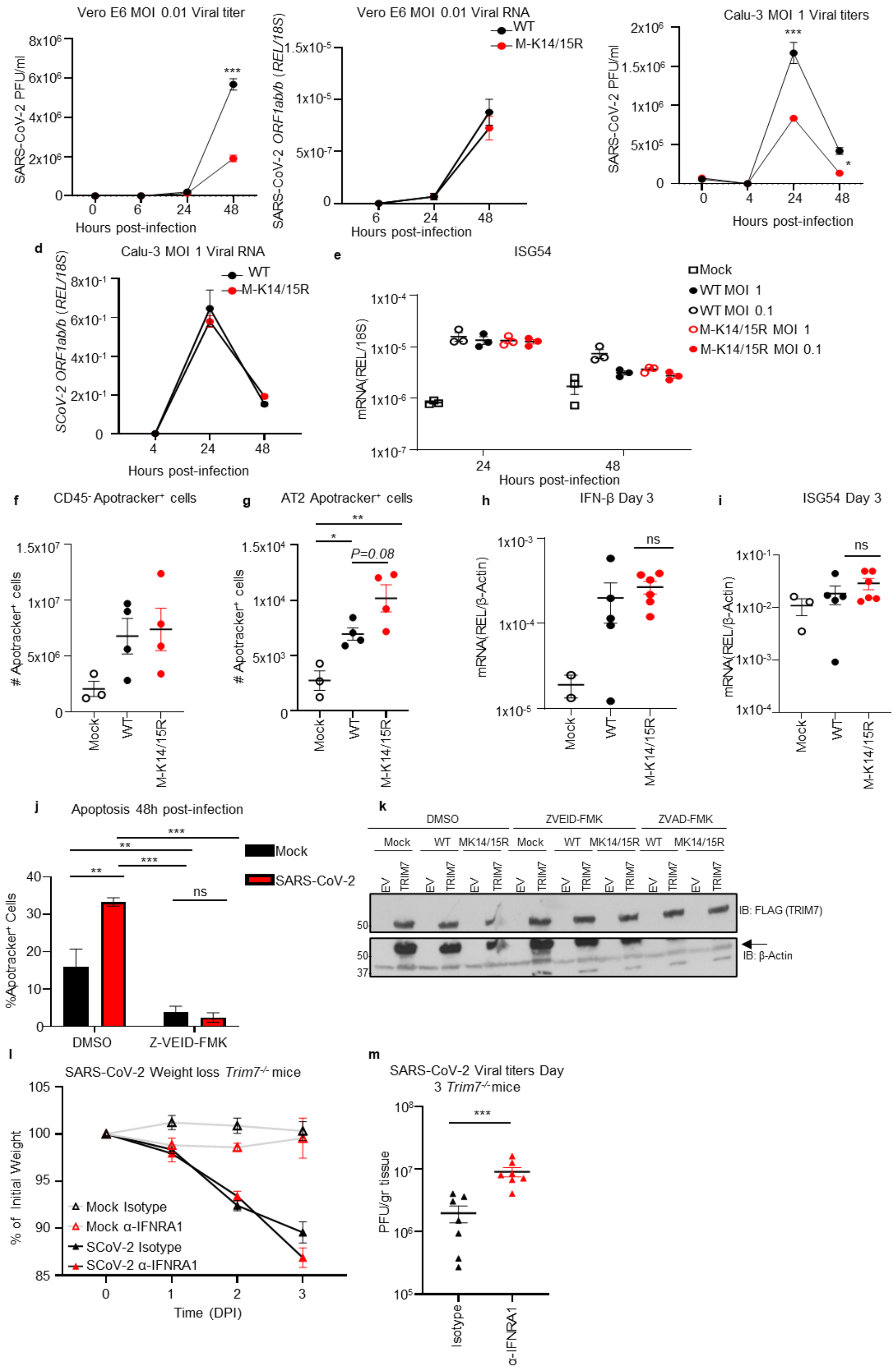


**Figure S6. M-K14/15R virus has defective budding but no IFN response is affected.** Vero E6 and Calu-3 infected with SARS-CoV-2 WT and M-K14/K15R MOI of 0.01 and 1 respectively. a) viral titers and b) viral RNA in Vero E6. c) viral titers and d) viral RNA in Calu-3. e) ISG54 expression in Calu-3 cells. WT mice infected with WT or M-K14/15R. f) lung $CD45^-$ cells $Apotracker^+$ and g) lung Alveolar Type 2 (AT2) cells ($CD45^-$ $CD31^-$ $CD24^-$ $Podoplanin^-$ $EpCAM^+$ $MHC\text{-}II^+$) positive for Apotracker staining. h) mRNA levels of IFN-β, and i) IS54. j) frequency of HEK 293T-hACE-2 cells infected with SARS-CoV-2 and treated with vehicle or 50μM of Z-VEID-FMK stained with Apotracker Green 48h post-infection. k) western blot analysis of expression of TRIM7-FLAG in 293T cells infected with CoV-2 WT and M-K14/K15R MOI 0.1 and treated with caspases inhibitors Z-VEID and Z-VAD. *Trim7*$^{-/-}$ mice treated with anti-IFNRA1 or isotype mock (n=3 males each group) at day 1 before infection and infected intranasal with SARS-CoV-2 CMA3p20 (n=7, 5 females and 2 males) for 3 days. l) weight loss and m) Viral lung titers. Data are depicted as Mean ± SEM. T-test analysis, one-way or 2-way Tukey's multiple comparisons tests. $p < 0.001$ **, $p < 0.0001$ ***, $p < 0.00001$ ****.

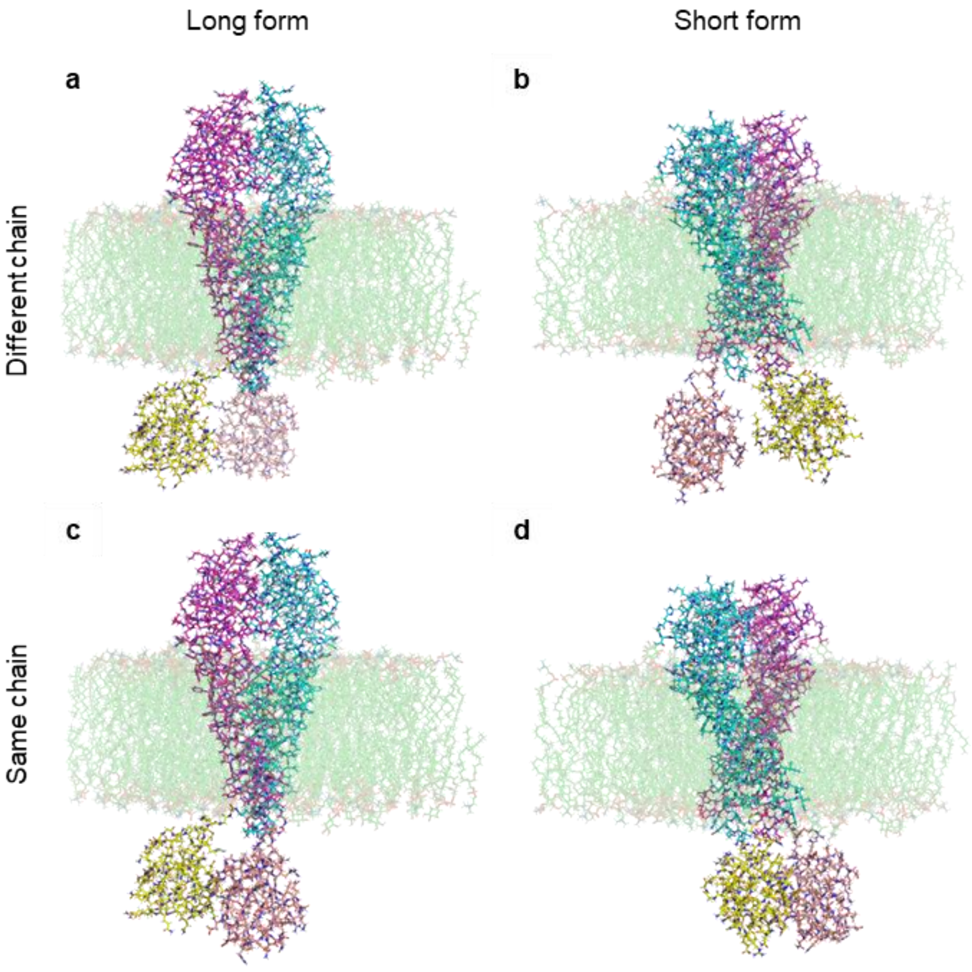


**Figure S7. Structural modeling of M. Modeling of different possible ubiquitinated M dimers**. The two chains of the M homodimer are shown in cyan and magenta whereas ubiquitin (Ub) is presented in yellow and salmon (the membrane is shown in green). a) long form of M (PDB 7VGR) with Ub attached to K14 and K15 in different chains. b) short form of M (PDB 7VGS) with Ub attached to K14 and K15 in different chains. c) long form of M with Ub attached to K14 and K15 in the same chain. d) short form of M with Ub attached to K14 and K15 of the same chain. Energetically based on the estimations with Surfaces, model A is the most favorable of the four, with full system ΔΔG calculations showing $\Delta\Delta G_{A\rightarrow B}$= 3.89 kcal/mol, $\Delta\Delta G_{A\rightarrow C}$= 1.01 kcal/mol, $\Delta\Delta G_{C\rightarrow D}$= 5.12 kcal/mol, and $\Delta\Delta G_{B\rightarrow D}$= 2.24 kcal/mol. In both cases (double ubiquitination in the same, or different chains), the long form is preferred.

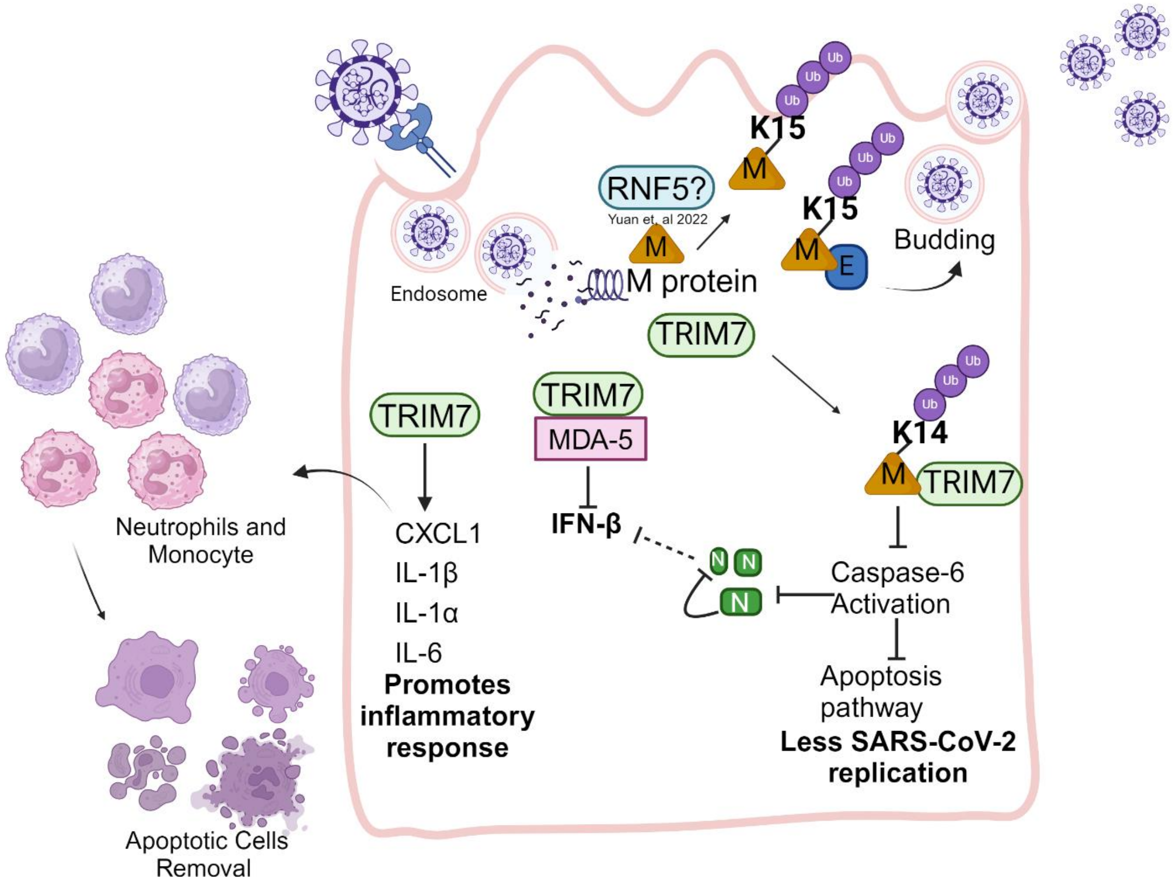


**Figure S8. Multifunctional role of TRIM7 during SARS-CoV-2 infection.** M protein can be ubiquitinated possibly by RNF5 on K15 residue, this is important for virus budding. On the other hand, TRIM7 can ubiquitinate M protein in the K14 residue, this ubiquitination is important to regulate caspase-6 activation and to inhibit apoptosis as well as to reduce the cleavage of N that could in turn inhibit the IFN-I production. TRIM7 can also interact with cytosolic receptor MDA-5 and inhibit the promoter activity of IFN-β by a mechanism that still unknown. TRIM7 is important to promote the production of pro-inflammatory cytokines IL-6, IL-1β and IL-1α as well as the chemokine CXCL1 that recruits neutrophils and monocytes to the lung, here these cells promote the removal of the apoptotic cells induced by SARS-CoV-2 and contribute with tissue repair. Created with BioRender.com.

Table 1. Mutations on Membrane protein K14 residue across clades.

| Clade | Residue Mutation | Sample Mutation count | Total Samples in Clade | Clade mutation % | Total Mutations | Clade Total Mutations % |
|---|---|---|---|---|---|---|
| 19A | K14del | 113 | 11833 | 0.955 | 113 | 0.955 |
| 19B | K14del | 11 | 7387 | 0.1489 | 11 | 0.149 |
| 20A | K14R | 3 | 120679 | 0.0025 | 274 | 0.227 |
| 20A | K14E | 1 | 120679 | 0.0008 | 274 | 0.227 |
| 20A | K14del | 270 | 120679 | 0.2237 | 274 | 0.227 |
| 20B | K14R | 3 | 104912 | 0.0029 | 137 | 0.131 |
| 20B | K14del | 134 | 104912 | 0.1277 | 137 | 0.131 |
| 21A | K14E | 2 | 53799 | 0.0037 | 2 | 0.004 |
| 20E | K14E | 4 | 103929 | 0.0038 | 8 | 0.008 |
| 20E | K14del | 4 | 103929 | 0.0038 | 8 | 0.008 |
| 20C | K14E | 8 | 68701 | 0.0116 | 54 | 0.079 |
| 20C | K14F | 1 | 68701 | 0.0015 | 54 | 0.079 |
| 20C | K14R | 1 | 68701 | 0.0015 | 54 | 0.079 |
| 20C | K14del | 44 | 68701 | 0.064 | 54 | 0.079 |
| 21H | K14E | 5 | 6144 | 0.0814 | 5 | 0.081 |
| 21B | K14R | 1 | 1045 | 0.0957 | 1 | 0.096 |
| 20I | K14* | 1 | 651108 | 0.0002 | 23 | 0.004 |
| 20I | K14R | 5 | 651108 | 0.0008 | 23 | 0.004 |
| 20I | K14Q | 1 | 651108 | 0.0002 | 23 | 0.004 |
| 20I | K14E | 5 | 651108 | 0.0008 | 23 | 0.004 |
| 20I | K14G | 1 | 651108 | 0.0002 | 23 | 0.004 |
| 20I | K14del | 7 | 651108 | 0.0011 | 23 | 0.004 |
| 20I | K14T | 3 | 651108 | 0.0005 | 23 | 0.004 |
| 20D | K14del | 9 | 6020 | 0.1495 | 9 | 0.15 |
| 21I | K14E | 2 | 151093 | 0.0013 | 7 | 0.005 |
| 21I | K14R | 1 | 151093 | 0.0007 | 7 | 0.005 |
| 21I | K14del | 4 | 151093 | 0.0026 | 7 | 0.005 |
| 21J | K14I | 8 | 2737780 | 0.0003 | 111 | 0.004 |
| 21J | K14R | 15 | 2737780 | 0.0005 | 111 | 0.004 |
| 21J | K14Q | 8 | 2737780 | 0.0003 | 111 | 0.004 |
| 21J | K14E | 60 | 2737780 | 0.0022 | 111 | 0.004 |
| 21J | K14del | 19 | 2737780 | 0.0007 | 111 | 0.004 |
| 21J | K14T | 1 | 2737780 | 0 | 111 | 0.004 |
| 20H | K14R | 14 | 9553 | 0.1466 | 16 | 0.167 |
| 20H | K14del | 2 | 9553 | 0.0209 | 16 | 0.167 |
| 21F | K14R | 1 | 33858 | 0.003 | 2 | 0.006 |
| 21F | K14del | 1 | 33858 | 0.003 | 2 | 0.006 |

| | | | | | | |
|---|---|---|---|---|---|---|
| 21C | K14E | 1 | 41628 | 0.0024 | 1 | 0.002 |
| 21K | K14E | 8 | 1591473 | 0.0005 | 70 | 0.004 |
| 21K | K14G | 1 | 1591473 | 0.0001 | 70 | 0.004 |
| 21K | K14del | 54 | 1591473 | 0.0034 | 70 | 0.004 |
| 21K | K14R | 7 | 1591473 | 0.0004 | 70 | 0.004 |
| 21L | K14E | 2 | 1143006 | 0.0002 | 57 | 0.005 |
| 21L | K14del | 46 | 1143006 | 0.004 | 57 | 0.005 |
| 21L | K14T | 5 | 1143006 | 0.0004 | 57 | 0.005 |
| 21L | K14R | 4 | 1143006 | 0.0003 | 57 | 0.005 |
| 22D | K14Q | 1 | 31187 | 0.0032 | 2 | 0.006 |
| 22D | K14R | 1 | 31187 | 0.0032 | 2 | 0.006 |
| 22F | K14R | 2 | 20443 | 0.0098 | 2 | 0.01 |
| 22C | K14R | 1 | 170929 | 0.0006 | 4 | 0.002 |
| 22C | K14E | 1 | 170929 | 0.0006 | 4 | 0.002 |
| 22C | K14G | 1 | 170929 | 0.0006 | 4 | 0.002 |
| 22C | K14del | 1 | 170929 | 0.0006 | 4 | 0.002 |
| 22B | K14E | 4 | 781430 | 0.0005 | 57 | 0.007 |
| 22B | K14del | 10 | 781430 | 0.0013 | 57 | 0.007 |
| 22B | K14R | 43 | 781430 | 0.0055 | 57 | 0.007 |
| 23C | K14R | 1 | 28556 | 0.0035 | 1 | 0.004 |
| 23A | K14E | 1 | 161212 | 0.0006 | 3 | 0.002 |
| 23A | K14R | 2 | 161212 | 0.0012 | 3 | 0.002 |
| 23B | K14R | 2 | 31125 | 0.0064 | 2 | 0.006 |
| 23E | K14E | 1 | 11253 | 0.0089 | 7 | 0.062 |
| 23E | K14R | 6 | 11253 | 0.0533 | 7 | 0.062 |
| 22E | K14R | 5 | 195894 | 0.0026 | 6 | 0.003 |
| 22E | K14del | 1 | 195894 | 0.0005 | 6 | 0.003 |

Table 2. Table 1. Mutations on Membrane protein K15 residue across clades.

| Clade | Residue Mutation | Sample Mutation count | Total Samples in Clade | Clade Individual mutation % | Total Mutations | Clade Total Mutation % |
|---|---|---|---|---|---|---|
| 19A | K15* | 3 | 11833 | 0.0254 | 4 | 0.034 |
| 19A | K15N | 1 | 11833 | 0.0085 | 4 | 0.034 |
| 20A | K15M | 2 | 120679 | 0.0017 | 19 | 0.016 |
| 20A | K15E | 8 | 120679 | 0.0066 | 19 | 0.016 |
| 20A | K15S | 3 | 120679 | 0.0025 | 19 | 0.016 |
| 20A | K15R | 1 | 120679 | 0.0008 | 19 | 0.016 |
| 20A | K15del | 5 | 120679 | 0.0041 | 19 | 0.016 |
| 19B | K15del | 1 | 7387 | 0.0135 | 35 | 0.474 |
| 19B | K15R | 34 | 7387 | 0.4603 | 35 | 0.474 |
| 20B | K15N | 26 | 104912 | 0.0248 | 32 | 0.031 |
| 20B | K15R | 2 | 104912 | 0.0019 | 32 | 0.031 |
| 20B | K15del | 4 | 104912 | 0.0038 | 32 | 0.031 |
| 21A | K15N | 7 | 53799 | 0.013 | 8 | 0.015 |
| 21A | K15E | 1 | 53799 | 0.0019 | 8 | 0.015 |
| 20C | K15N | 55 | 68701 | 0.0801 | 106 | 0.154 |
| 20C | K15R | 50 | 68701 | 0.0728 | 106 | 0.154 |
| 20C | K15E | 1 | 68701 | 0.0015 | 106 | 0.154 |
| 21H | K15N | 1 | 6144 | 0.0163 | 1 | 0.016 |
| 20E | K15N | 19 | 103929 | 0.0183 | 25 | 0.024 |
| 20E | K15T | 1 | 103929 | 0.001 | 25 | 0.024 |
| 20E | K15R | 2 | 103929 | 0.0019 | 25 | 0.024 |
| 20E | K15del | 3 | 103929 | 0.0029 | 25 | 0.024 |
| 20J | K15Q | 2 | 29088 | 0.0069 | 2 | 0.007 |
| 20I | K15M | 7 | 651108 | 0.0011 | 102 | 0.016 |
| 20I | K15N | 22 | 651108 | 0.0034 | 102 | 0.016 |
| 20I | K15E | 26 | 651108 | 0.004 | 102 | 0.016 |
| 20I | K15T | 1 | 651108 | 0.0002 | 102 | 0.016 |
| 20I | K15R | 40 | 651108 | 0.0061 | 102 | 0.016 |
| 20I | K15del | 6 | 651108 | 0.0009 | 102 | 0.016 |
| 20D | K15N | 1 | 6020 | 0.0166 | 1 | 0.017 |
| 20D | K15del | 1 | 6020 | 0.0166 | 1 | 0.017 |
| 21I | K15N | 4 | 151093 | 0.0026 | 37 | 0.024 |
| 21I | K15Q | 1 | 151093 | 0.0007 | 37 | 0.024 |
| 21I | K15E | 24 | 151093 | 0.0159 | 37 | 0.024 |
| 21I | K15del | 3 | 151093 | 0.002 | 37 | 0.024 |
| 21I | K15R | 5 | 151093 | 0.0033 | 37 | 0.024 |
| 21J | K15N | 2433 | 2737780 | 0.0889 | 2785 | 0.102 |
| 21J | K15Q | 165 | 2737780 | 0.006 | 2785 | 0.102 |
| 21J | K15T | 7 | 2737780 | 0.0003 | 2785 | 0.102 |
| 21J | K15R | 123 | 2737780 | 0.0045 | 2785 | 0.102 |
| 21J | K15del | 29 | 2737780 | 0.0011 | 2785 | 0.102 |
| 21J | K15M | 11 | 2737780 | 0.0004 | 2785 | 0.102 |

| | | | | | | |
|---|---|---|---|---|---|---|
| 21J | K15E | 17 | 2737780 | 0.0006 | 2785 | 0.102 |
| 20H | K15del | 2 | 9553 | 0.0209 | 3 | 0.031 |
| 20H | K15R | 1 | 9553 | 0.0105 | 3 | 0.031 |
| 21F | K15del | 1 | 33858 | 0.003 | 1 | 0.003 |
| 21C | K15N | 1 | 41628 | 0.0024 | 1 | 0.002 |
| 20G | K15N | 4 | 83603 | 0.0048 | 6 | 0.007 |
| 20G | K15R | 2 | 83603 | 0.0024 | 6 | 0.007 |
| 21K | K15T | 1 | 1591473 | 0.0001 | 166 | 0.01 |
| 21K | K15N | 31 | 1591473 | 0.0019 | 166 | 0.01 |
| 21K | K15R | 35 | 1591473 | 0.0022 | 166 | 0.01 |
| 21K | K15Q | 22 | 1591473 | 0.0014 | 166 | 0.01 |
| 21K | K15del | 55 | 1591473 | 0.0035 | 166 | 0.01 |
| 21K | K15M | 6 | 1591473 | 0.0004 | 166 | 0.01 |
| 21K | K15E | 16 | 1591473 | 0.001 | 166 | 0.01 |
| 21L | K15N | 7 | 1143006 | 0.0006 | 71 | 0.006 |
| 21L | K15R | 11 | 1143006 | 0.001 | 71 | 0.006 |
| 21L | K15del | 52 | 1143006 | 0.0045 | 71 | 0.006 |
| 21L | K15E | 1 | 1143006 | 0.0001 | 71 | 0.006 |
| 22D | K15N | 5 | 31187 | 0.016 | 7 | 0.022 |
| 22D | K15R | 2 | 31187 | 0.0064 | 7 | 0.022 |
| 22F | K15del | 1 | 20443 | 0.0049 | 5 | 0.024 |
| 22F | K15N | 1 | 20443 | 0.0049 | 5 | 0.024 |
| 22F | K15R | 3 | 20443 | 0.0147 | 5 | 0.024 |
| 22C | K15N | 1 | 170929 | 0.0006 | 7 | 0.004 |
| 22C | K15R | 5 | 170929 | 0.0029 | 7 | 0.004 |
| 22C | K15del | 1 | 170929 | 0.0006 | 7 | 0.004 |
| 22B | K15del | 13 | 781430 | 0.0017 | 106 | 0.014 |
| 22B | K15N | 6 | 781430 | 0.0008 | 106 | 0.014 |
| 22B | K15E | 3 | 781430 | 0.0004 | 106 | 0.014 |
| 22B | K15R | 79 | 781430 | 0.0101 | 106 | 0.014 |
| 22B | K15Q | 5 | 781430 | 0.0006 | 106 | 0.014 |
| 22A | K15N | 1 | 95899 | 0.001 | 1 | 0.001 |
| 23C | K15R | 1 | 28556 | 0.0035 | 1 | 0.004 |
| 23A | K15N | 4 | 161212 | 0.0025 | 11 | 0.007 |
| 23A | K15R | 7 | 161212 | 0.0043 | 11 | 0.007 |
| 23B | K15del | 1 | 31125 | 0.0032 | 1 | 0.003 |
| 23D | K15N | 69 | 36429 | 0.1894 | 72 | 0.198 |
| 23D | K15Q | 1 | 36429 | 0.0027 | 72 | 0.198 |
| 23D | K15R | 1 | 36429 | 0.0027 | 72 | 0.198 |
| 23D | K15E | 1 | 36429 | 0.0027 | 72 | 0.198 |
| 23E | K15R | 7 | 11253 | 0.0622 | 7 | 0.062 |
| 22E | K15del | 1 | 195894 | 0.0005 | 4 | 0.002 |
| 22E | K15M | 1 | 195894 | 0.0005 | 4 | 0.002 |
| 22E | K15N | 1 | 195894 | 0.0005 | 4 | 0.002 |
| 22E | K15R | 1 | 195894 | 0.0005 | 4 | 0.002 |
| 23F | K15N | 1 | 31861 | 0.0031 | 4 | 0.013 |

| | | | | | | |
|---|---|---|---|---|---|---|
| 23F | K15Q | 2 | 31861 | 0.0063 | 4 | 0.013 |
| 23F | K15R | 1 | 31861 | 0.0031 | 4 | 0.013 |

Table 3. Table 1. Mutations on Membrane protein K14/K15 residue across clades.

| Clade | Residue Mutation | Sample Mutation count | Total Samples in Clade | Clade mutation % | Total Mutations | Clade Total Mutations % |
|---|---|---|---|---|---|---|
| 19B | K14del, K15del | 1 | 11833 | 0.008 | 1 | 0.008 |
| 20A | K14del, K15del | 4 | 120679 | 0.003 | 4 | 0.003 |
| 20B | K14del, K15del | 2 | 104912 | 0.002 | 2 | 0.002 |
| 20I | K14del, K15del | 6 | 651108 | 0.001 | 6 | 0.001 |
| 20D | K14del, K15del | 1 | 6020 | 0.017 | 1 | 0.017 |
| 20E | K14del, K15del | 3 | 103929 | 0.003 | 3 | 0.003 |
| 21I | K14del, K15del | 3 | 151093 | 0.002 | 3 | 0.002 |
| 21J | K14del, K15del | 19 | 2737780 | 0.001 | 19 | 0.001 |
| 20H | K14del, K15del | 2 | 9553 | 0.021 | 2 | 0.021 |
| 21F | K14del, K15del | 1 | 33858 | 0.003 | 1 | 0.003 |
| 21K | K14del, K15del | 51 | 1591473 | 0.003 | 52 | 0.003 |
| 21K | K14R, K15Q | 1 | 1591473 | 0 | 52 | 0.003 |
| 21L | K14del, K15del | 45 | 1143006 | 0.004 | 45 | 0.004 |
| 22C | K14del, K15del | 2 | 170929 | 0.001 | 2 | 0.001 |
| 22B | K14del, K15del | 11 | 781430 | 0.001 | 11 | 0.001 |
| 22E | K14del, K15del | 1 | 195894 | 0.001 | 1 | 0.001 |
